# Genome-wide CRISPR screens of T cell exhaustion identify chromatin remodeling factors that limit T cell persistence

**DOI:** 10.1101/2022.04.20.488974

**Authors:** Julia A. Belk, Winnie Yao, Nghi Ly, Katherine A. Freitas, Yan-Ting Chen, Quanming Shi, Alfredo M. Valencia, Eric Shifrut, Nupura Kale, Kathryn E. Yost, Connor V. Duffy, Madeline A. Hwee, Zhuang Miao, Alan Ashworth, Crystal L. Mackall, Alexander Marson, Julia Carnevale, Santosh A. Vardhana, Ansuman T. Satpathy

## Abstract

T cell exhaustion limits anti-tumor immunity, but the molecular determinants of this process remain poorly understood. Using a chronic antigen stimulation assay, we performed genome-wide CRISPR/Cas9 screens to systematically discover genetic regulators of T cell exhaustion, which identified an enrichment of epigenetic factors. *In vivo* CRISPR screens in murine and human tumor models demonstrated that perturbation of several epigenetic regulators, including members of the INO80 and BAF chromatin remodeling complexes, improved T cell persistence in tumors. *In vivo* paired CRISPR perturbation and single-cell RNA sequencing revealed distinct transcriptional roles of each complex and that depletion of canonical BAF complex members, including *Arid1a*, resulted in the maintenance of an effector program and downregulation of terminal exhaustion-related genes in tumor-infiltrating T cells. Finally, *Arid1a*-depletion limited the global acquisition of chromatin accessibility associated with T cell exhaustion and led to improved anti-tumor immunity after adoptive cell therapy. In summary, we provide a comprehensive atlas of the genetic regulators of T cell exhaustion and demonstrate that modulation of the epigenetic state of T cell exhaustion can improve T cell responses in cancer immunotherapy.

## Main Text

T cell exhaustion is a process that is driven by chronic T cell receptor (TCR) stimulation and induces the stable expression of inhibitory surface receptors, poor response to tumor antigens, and low cell proliferation and persistence *in vivo* (Wherry and Kurachi, 2015; Collier et al., 2021). Originally identified in the setting of chronic viral infection (Zajac et al., 1998; Barber et al., 2006), T cell exhaustion is now appreciated to occur in diverse disease settings, including cancer and autoimmune disease (McKinney et al., 2015; McLane et al., 2019). Importantly, studies have demonstrated that T cell exhaustion represents a major barrier for the efficacy of both checkpoint blockade and chimeric antigen receptor T (CAR-T) cell immunotherapies, and that manipulating this process may lead to improved efficacy of T cell responses in cancer (Sakuishi et al., 2010; Long et al., 2015; Fraietta et al., 2018a, 2018b; Ribas and Wolchok, 2018; Lynn et al., 2019; Yost et al., 2019; Weber et al., 2021).

Until recently, our understanding of the molecular determinants of this process was limited, and the prevailing view was that T cell exhaustion may largely reflect the aberrant expression of a small set of dysfunctional genes (e.g. PD-1), rather than a unique cell state or differentiation pathway. However, recent genomic studies in murine models of chronic infection and cancer have demonstrated that T cell exhaustion is associated with a broad remodeling of the transcriptional and epigenomic landscape, which is conserved across disease settings (Pauken et al., 2016; Sen et al., 2016; Philip et al., 2017; Pritykin et al., 2021). This unique epigenetic state is primarily driven by chronic antigen and TCR signaling, and results in a stable cellular phenotype that is not changed by anti-PD-1 treatment (Pauken et al., 2016; Pritykin et al., 2021; Schietinger et al., 2016). Indeed, in cancer patients receiving PD-1 blockade, exhausted T cells display a distinct differentiation trajectory and end-stage chromatin profile, compared to functional effector T cells, and clonal tracing of exhausted T cells demonstrated that these cells are limited in their capacity to proliferate and perform effector functions in response to immunotherapy (Yost et al., 2019; Philip et al., 2017; Satpathy et al., 2019).

CRISPR/Cas9 screening has emerged as a powerful discovery tool for the molecular determinants of immune cell differentiation and function (Parnas et al., 2015; Shalem et al., 2014; Shifrut et al., 2018; Wang et al., 2014). For example, prior CRISPR/Cas9 screens in T cells have been used to identify transcription factors and metabolic regulators of T cell fate *in vivo*, as well as therapeutic targets (Chen et al., 2021; Dong et al., 2019; Huang et al., 2021; LaFleur et al., 2019; Wei et al., 2019). However, inherent limitations in scaling these *in vivo* assays have constrained library diversity of these screens, largely preventing genome-wide analysis and an unbiased discovery of novel regulators of T cell phenotypes. Furthermore, assays that simultaneously screen for multiple functions of T cells—for example, tissue localization, infiltration, and differentiation in tumors—have also made it challenging to interrogate the impact of a particular gene perturbation on a single aspect of T cell function and phenotype, such as exhaustion.

Here, we developed an *in vitro* model of CD8^+^ T cell exhaustion, which recapitulates the epigenomic features of exhaustion that are observed *in vivo* and is scalable for genome-wide CRISPR/Cas9 screens. Using this model, we provide a systematic and comprehensive view of the genetic regulators of T cell exhaustion and identify dozens of novel factors which have not previously been implicated in this process. Strikingly, these factors are enriched for chromatin remodeling proteins, including subunits of the INO80 (inositol requiring mutant 80) nucleosome positioning complex and the SWI/SNF (switch/sucrose non-fermentable) chromatin remodeling complex. *In vivo* CRISPR screens and T cell transfer experiments revealed that depletion of INO80 and canonical BRG1 or BRM-associated factor (cBAF; SWI/SNF family) complex members—in particular, *Arid1a*—led to increased persistence of T cells. *In vivo* Perturb-seq analysis revealed that depletion of these factors resulted in the upregulation of an effector transcriptional program in exhausted T cells and that each complex regulated a distinct set of target genes; depletion of INO80 factors led to the upregulation of metabolic gene expression, while depletion of cBAF factors led to the upregulation of effector molecule gene expression and the downregulation of exhaustion-related genes. ATAC-seq profiling of *Arid1a*-depleted T cells demonstrated that *Arid1a* was required for the acquisition of exhaustion-associated chromatin remodeling that occurs during chronic antigen stimulation. Finally, *Arid1a*-depleted cells exhibited improved tumor control, suggesting that modulation of the epigenetic state of T cell exhaustion via chromatin remodeling factors may be an effective path to improve T cell responses in cancer immunotherapy.

## Results

### An *in vitro* chronic stimulation assay recapitulates the epigenetic program of terminal T cell exhaustion

To develop an assay that is amenable to genome-wide CRISPR/Cas9 screening of T cell exhaustion, we adapted our previously described approach, which used anti-CD3 antibodies to enforce clustering of the T cell co-receptor, CD3, and thereby induce chronic TCR signaling in an antigen-independent manner (Figure 1A) (Vardhana et al., 2020). Compared to *in vivo* assays, this model isolates the core determinant of T cell exhaustion—chronic stimulation through the TCR complex—and removes T cell localization and trafficking effects, as well as immunosuppressive factors in the tumor microenvironment (TME). Importantly, this assay is scalable; we were able to culture upwards of 10^8^ cells, enabling coverage of genome-wide CRISPR sgRNA libraries. Over the course of 8 days of anti-CD3 stimulation (after 2 days of anti-CD3/CD28 activation), we confirmed a progressive upregulation of the inhibitory receptors, PD-1 and TIM3, and a growth defect in the chronically-stimulated T cells, compared to cells passaged without further TCR stimulation after initial activation (acute stimulation; p < 0.0001, unpaired t-test; Figure 1B-C, S1A). Chronically stimulated T cells also exhibited defects in the secretion of IFNɣ and TNFɑ after restimulation with phorbol myristate acetate and ionomycin, compared to acutely stimulated cells (Figure S1B). Coculture of OT-1 T cells and B16 tumor cells expressing Luciferase and pulsed with the cognate peptide antigen, SIINFEKL, also demonstrated that chronically stimulated cells were impaired in tumor killing *in vitro* (Figure S1C). Finally, transplant of chronically stimulated OT-1 T cells into mice bearing B16-OVA tumors demonstrated reduced tumor control *in vivo*, compared to transplant of acutely stimulated T cells (average tumor size 20 days after transplant: 1,849.6 mm^3^ (chronic) or 755.0 mm^3^ (acute); p = 0.005, unpaired t-test; Figure S1D).

**Figure 1:**
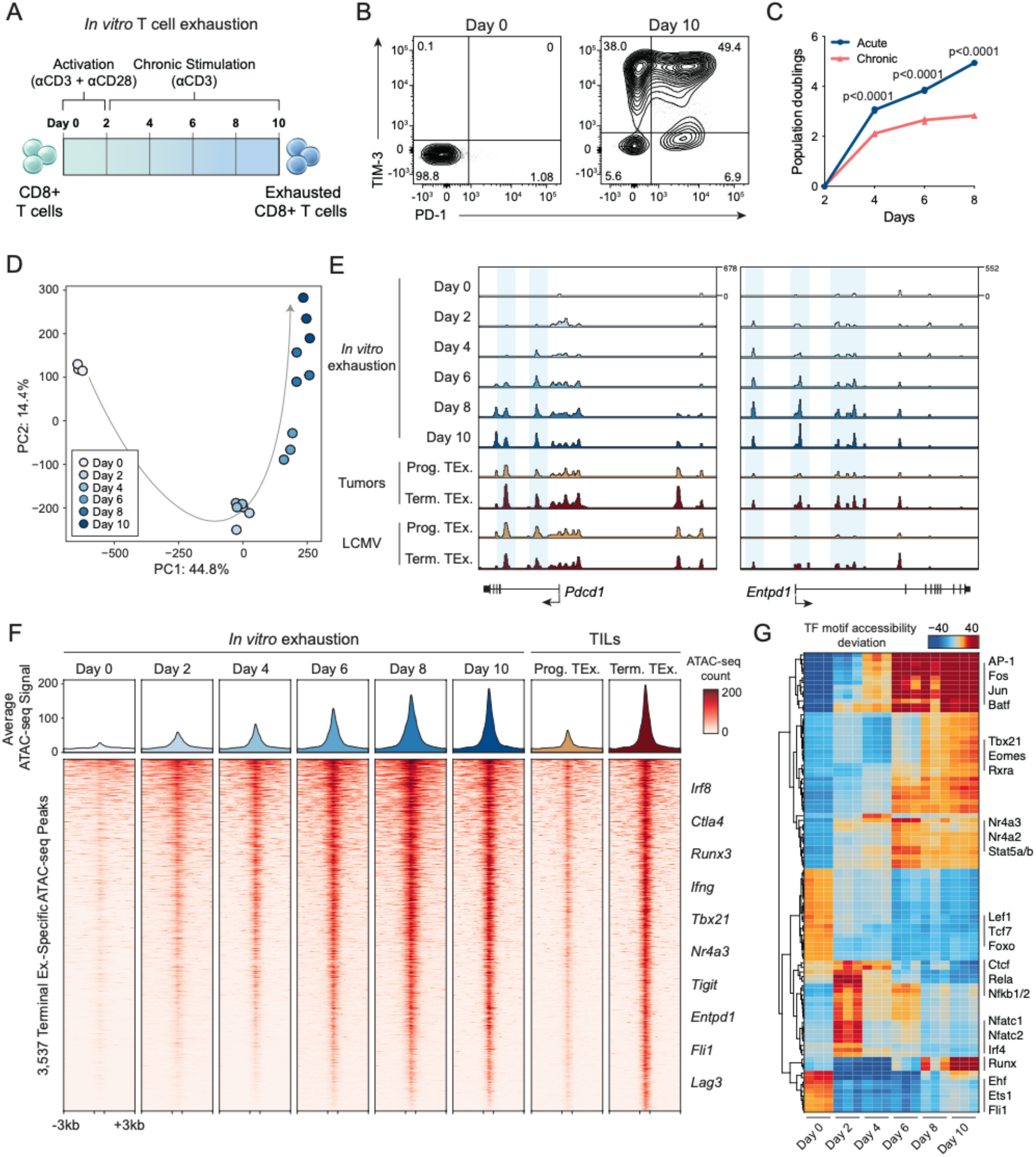
*In vitro* chronic antigen stimulation assay recapitulates the epigenetic hallmarks of T cell exhaustion. **(A)** Diagram of *in vitro* exhaustion assay. **(B)** Surface phenotype of CD8^+^ T cells at day 0 and day 10 of the T cell exhaustion assay, gated on live cells. **(C)** Expansion of chronically stimulated and acutely stimulated T cells *in vitro*. **(D)** Principal component analysis of ATAC-seq profiles of CD8^+^ T cells throughout the course of chronic stimulation. **(E)** ATAC-seq signal tracks in the *Pdcd1* and *Entpd1* gene loci at each time point in the *in vitro* exhaustion assay, as well as previously published reference ATAC-seq profiles from T cells in tumors or LCMV (Miller et al., 2019). **(F)** Heatmap showing ATAC-seq coverage of each peak in the “Terminal Exhaustion peak set” for each time point in the *in vitro* exhaustion assay. Reference data from TILs is also included. Selected nearest genes are indicated on the right. **(G)** chromVAR motif accessibility heatmap for each ATAC-seq sample. Selected motifs are indicated on the right. Top 100 most variable motifs are shown.

We next asked whether the *in vitro* exhaustion assay recapitulated epigenetic hallmarks of T cell exhaustion *in vivo* (Pauken et al., 2016; Sen et al., 2016; Satpathy et al., 2019). We performed the assay for transposase-accessible chromatin with sequencing (ATAC-seq) over the course of chronic stimulation and analyzed global chromatin accessibility profiles. Principal component analysis (PCA) of ATAC-seq profiles showed that PC1 separated naïve cells (Day 0) from all other samples, while PC2 captured a progressive epigenetic differentiation of the T cells during chronic stimulation (Figure 1D). Analysis of individual gene loci, including *Pdcd1* and *Entpd1,* demonstrated an increase in accessibility at known exhaustion-specific regulatory elements (Figure 1E) (Miller et al., 2019). We evaluated the global epigenetic similarity of *in vitro* stimulated cells to reference T cell exhaustion data from tumors and chronic infection (Miller et al., 2019). We defined a “terminal exhaustion peak set” as ATAC-seq peaks that are specifically active in terminally exhausted T cells, compared to progenitor exhausted T cells, and we identified 3,537 terminal exhaustion ATAC-seq peaks in the B16 melanoma tumor model and 2,346 peaks in the lymphocytic choriomeningitis virus (LCMV) chronic infection model (Log_2_ FC ≥ 1; FDR ≤ 0.05; Figure 1F, S1E). A comparison of terminal exhaustion peak accessibility in each model with the *in vitro* exhaustion ATAC-seq data demonstrated that the *in vitro* assay recapitulated global epigenomic changes observed *in vivo*: 88.6% of ATAC-seq peaks in tumors and 70.1% of ATAC-seq peaks in chronic infection showed a shared increase in accessibility in the *in vitro* model at Day 10 (FDR ≤ 0.05; Figure 1F, S1E). Finally, we analyzed chromatin accessibility at transcription factor (TF) binding sites using chromVAR (Schep et al., 2017), which showed that TF motifs previously associated with terminal exhaustion, including Batf, Fos, Jun, and Nr4a motifs were highly accessible *in vitro* at day 10. Moreover, we observed progressive loss of accessibility at naive and progenitor exhaustion-associated Lef1 and Tcf7 motifs, early increased accessibility of NF-κB and Nfat motifs, and later increased accessibility of AP-1 and Nr4a motifs, mirroring the progression of TF activity observed in T cell exhaustion *in vivo* (Figure 1G) (Lynn et al., 2019; Miller et al., 2019; Beltra et al., 2020; Daniel et al., 2021). In summary, these results demonstrate that the *in vitro* T cell exhaustion assay displayed hallmark functional and genomic features of *in vivo* T cell exhaustion, including expression of inhibitory receptors, impaired proliferation, cytokine secretion, and tumor killing, and global chromatin remodeling of dysfunctional T cell gene loci.

### Genome-wide CRISPR screens identify genetic regulators of T cell exhaustion

We next adapted the *in vitro* exhaustion assay to be compatible with CRISPR screening (Figure 2A). We isolated CD8^+^ T cells from Rosa26-Cas9 knockin mice, which constitutively express Cas9-P2A-EGFP (Platt et al., 2014). 24 hours after T cell isolation, we transduced Cas9^+^CD8^+^ T cells with a genome-wide retroviral sgRNA library containing 90,230 sgRNAs (Henriksson et al., 2019). We introduced a 48-hour delay between activation and the onset of chronic stimulation to allow time for efficient gene editing and puromycin selection of transduced cells. To identify genes that specifically modulated fitness in the presence of chronic stimulation, we split the cells into acute (IL-2 only) and chronic (anti-CD3 and IL-2) stimulation conditions on Day 4 and sequenced both pools on Day 10 (Figure 2A). We performed replicate screens and confirmed: (1) transduction of T cells at low multiplicity of infection (MOI) to optimize single sgRNA targeting of cells (replicate 1: 16.9% sgRNA^+^ cells, MOI = 0.18; replicate 2: 29.3% sgRNA^+^ cells, MOI = 0.35; Figure S2A), and (2) T cell exhaustion in this CRISPR-adapted assay by analyzing cell surface phenotype on Day 10 of the chronic culture (Figure S2B). Next, we analyzed the guide representation in each condition: of the 90,230 sgRNAs present in the plasmid library design, >99% were recovered in each acute sample (acute replicate 1: 89,329 (99.0%); acute replicate 2: 89,625 (99.3%); Figure S2C, **Table S1**). The “chronic” samples demonstrated more evidence of selective pressure, with a wider spread in guide counts and a greater number of guides dropping out of the screen (chronic replicate 1: 75,776 sgRNAs detected (83.9%); chronic replicate 2: 87,524 sgRNAs detected (97.0%); Figure S2C, **Table S1**). Comparing the counts observed for each sgRNA in the acute and chronic conditions revealed a positive correlation, as expected, with a small population of sgRNAs dramatically enriched in the chronic condition (Figure S2D).

**Figure 2:**
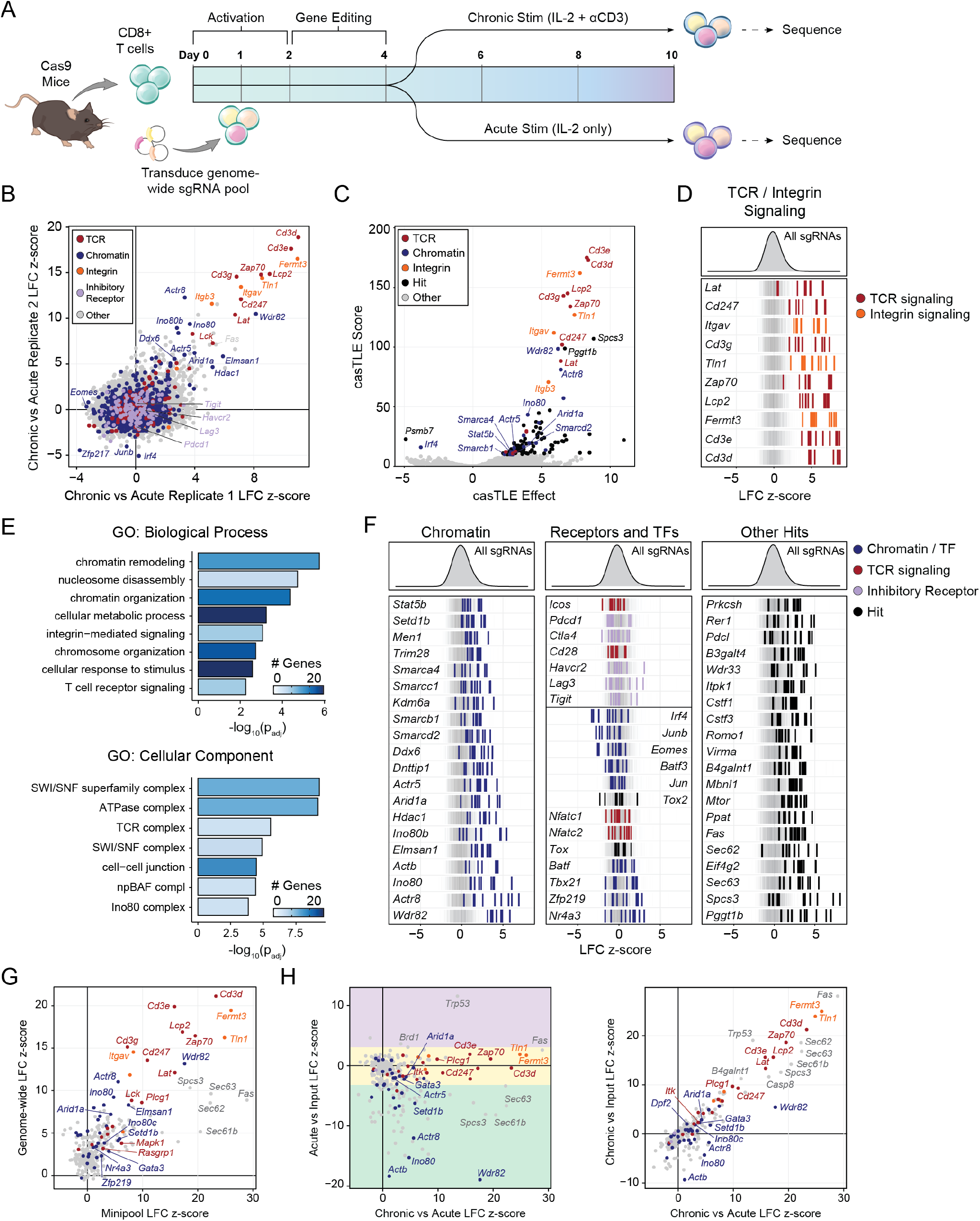
Genome-wide functional interrogation of T cell exhaustion. **(A)** Diagram of genome-wide T cell exhaustion screen. **(B)** Correlation of replicate screens with selected functional categories of genes colored as indicated. Gene sets were based on GO Terms and were supplemented with manual annotations. **(C)** casTLE volcano plot of the Chronic vs Acute stimulation screen comparison, with top hits labeled. **(D)** Individual sgRNA z-scores for top hits in “integrin signaling” or “TCR signaling” functional categories. **(E)** GO Term analysis of the top 100 positive hits. **(F)** Individual sgRNA z-scores for genes in different functional categories: chromatin (left), selected receptors and transcription factors (center), or other (right). Histogram for all guide z-scores is shown above, and 1,000 randomly selected guides are shown in the background of each row in grey, for visual reference. **(G)** Correlation of Acute vs Chronic z-scores in the mini-pool versus the genome-wide screen. **(H)** Correlation of the mini-pool Chronic vs Acute z-scores against Acute vs Input (left) or Chronic vs Input (right). Genes in (G) and (H) are colored by functional category: TCR signaling (red), integrin signaling (orange), chromatin (blue), or other (grey). Colored boxes in (H, left) denote enhanced (purple), similar (yellow), or reduced (green) expansion after acute stimulation.

Positive controls for the screen are components of the TCR signaling pathway, since depletion of these factors impairs antigen-driven (or anti-CD3-driven) signaling, and therefore, mitigates exhaustion in this model. Accordingly, we first analyzed the enrichments of the CD3 receptor subunits (*Cd3e*, *Cd3d*, *Cd3g*, *Cd247*; Figure S2D in red) and observed a robust enrichment of guides targeting these genes in both replicates. We next normalized the counts table, computed z-scores for each sgRNA, and merged these sgRNA-level z-scores into a z-score for each gene, as previously described (Flynn et al., 2021). Merging the replicates yielded an overall z-score and ranking for each gene (Figure 2B-C, **Table S2**). We validated this analysis approach by comparing screen hits obtained from two additional widely adopted CRISPR sgRNA enrichment analytical methods, MAGeCK and casTLE, which showed a high correlation between effect size estimates (casTLE effect size correlation: R = 0.66; MAGeCK log fold change correlation: R = 0.77; Figure S3A-D, **Table S2**) (Li et al., 2014; Morgens et al., 2016). A comparison of the genes classified as hits using each method revealed that the largest group of hits were shared by all three methods (“hit” corresponds to FDR < 0.05 for our pipeline and MAGeCK or casTLE score > 10; 48 genes; Figure S3B). Finally, we sought to ensure that the identification of screen hits was robust to the choice of reference sgRNAs. The retroviral library pool did not contain a control sgRNA set, so we compared our normalization strategy (relative to all sgRNAs in the pool) to a strategy that utilizes a set of sgRNAs targeting olfactory receptors that are not expressed or predicted to function in T cells (Gilbert et al., 2014). We found that normalizing sgRNA enrichments to the olfactory receptor sgRNA set modestly boosted the statistical power of the screen results but otherwise had a minimal impact on the results (Figure S3E).

Enriched genes are those that restrict T cell persistence in the presence of chronic antigen (deletion improves persistence), while depleted genes represent “essential genes” for persistence. In addition to *Cd3e*, *Cd3d,* and *Cd3g,* top ranked genes in the screen included other known components of the TCR signaling pathway, such as *Zap70*, *Lcp2*, *Lat*, and *Lck*, as well as cell adhesion and integrin-related genes *Fermt3*, *Tln1*, *Itgav*, and *Itgb3* (Figure 2B-D). GO Term analysis of the top 100 positive regulators of exhaustion confirmed that the “T cell receptor signaling pathway” term was highly enriched (p_adj_ = 5.44 x 10^-3^; Figure 2E). Surprisingly, in addition to TCR-related GO terms, the other top terms were related to epigenetics, including “chromatin remodeling” (p_adj_ = 1.74 x 10^-6^), “nucleosome disassembly” (p_adj_ = 1.90 x 10^-5^), and “chromatin organization” (p_adj_ = 4.14 x 10^-5^; Figure 2E). Indeed, analysis of additional highly enriched genes identified a number of chromatin-related factors, including *Arid1a, Smarcc1, Smarcd2, Ino80, Actr8,* and *Actr5* (Figure 2F**, left**). Of note, the co-stimulatory and inhibitory receptors *Icos*, *Pdcd1*, *Ctla4*, *Cd28*, *Havcr2*, *Lag3*, and *Tigit* were not significantly enriched by the screen (Figure 2F**, center**), perhaps supporting evidence that inhibitory receptor signaling does not directly impact the process of T cell exhaustion or improve the persistence of engineered T cells in humans (Pauken et al., 2016; Weber et al., 2021; Yost et al., 2019; Stadtmauer et al., 2020; Yost et al., 2021; Wei et al., 2017). Among TFs, *Irf4*, *Junb*, *Eomes*, and *Batf3* were depleted, while *Tbx21* and *Nr4a3* were modestly enriched, supporting previous demonstrations of their roles in exhaustion (Figure 2F**, center**) (Ataide et al., 2020; Chen et al., 2019; Paley et al., 2012; Seo et al., 2021). In contrast, *Tox* and *Tox2*, which are critical for the development of exhaustion, were not hits, supporting previous studies demonstrating that deletion of these factors may not improve T cell persistence *in vivo*, perhaps due to activation-induced cell death (Figure 2F**, center**) (Alfei et al., 2019; Khan et al., 2019; Scott et al., 2019). Similarly, *Jun* and *Batf* were not hits, suggesting that while overexpression of these factors improves T cell persistence, deletion does not (Lynn et al., 2019; Seo et al., 2021).

We next used Cytoscape to visualize the protein-protein interaction network of top enriched and depleted genes (Figure S4) (Shannon et al., 2003). This analysis confirmed the highly interconnected and enriched network of factors that directly associate with the TCR complex and downstream signaling components, as well as several other protein complexes and functional categories. These included the INO80 nucleosome positioning complex (hits included *Ino80*, *Ino80b*, *Actr5*, and *Actr8*), the Set1C/COMPASS complex that regulates histone methylation (hits included *Wdr82*, *Dpy30*, and *Setd1b*), the BAF chromatin remodeling complex (hits included *Arid1a*, *Smarcb1*, *Smarcd2*, *Smarca4*, and *Smarcc1*), and the mitotic deacetylase (MiDAC) complex comprised of *Hdac1*, *Dnttip1*, and *Elmsan1* (Mondal et al., 2020). Within the BAF complex subunits, *Smarcc1*, *Smarcd2*, and *Smarcb1* are part of the BAF core, which assembles together with *Arid1a*, *Smarca4* (an ATPase), *Actb*, and other components to form the BAF complex (Mashtalir et al., 2018). We also observed enrichment of the mRNA-processing CSTF complex (*Cstf1*, *Cstf2*, and *Cstf3*), N^6^-methyladenosine (m^6^A) RNA modification-related genes, (*Zfp217*, *Rbm15*, and *Virma*), and several hits related to the endoplasmic reticulum and protein secretion (*Spcs2*, *Spcs3*, *Sec62*, *Sec63*), lipid biosynthesis (*Gpi1*, *Pigv*, *Dpm3*), and the Mitochondrial Complex V (*Atp5b*, *Atp5d*, *Atp5a1*).

Finally, to understand the expression of these genes in the context of exhausted T cell subsets *in vivo*, we analyzed their expression patterns in previous single-cell RNA-seq data of exhausted T cells in chronic viral infection (Figure S5A-B) (Raju et al., 2021). This dataset encompasses the key subtypes of progenitor, transitory, and terminally exhausted T cells (Hudson et al., 2019; Zander et al., 2019), and analysis of the top 100 ranked genes demonstrated that all factors are detectably expressed in T cells in chronic viral infection (Figure S5C-D). Moreover, all but two of these genes, *Tmem253* and *Itgb3*, were expressed early during exhaustion (98 of the top 100 hits detectably expressed in progenitor exhausted T cells) and remained relatively stable across exhausted subtypes, suggesting that disruption of epigenetic factors and other novel hits may act early during the cellular course of T cell exhaustion, rather than reversing exhaustion after terminal differentiation (Figure S5C). In summary, the *in vitro* genome-wide CRISPR/Cas9 exhaustion screen provides an unbiased and comprehensive catalog of genetic factors that govern the process of chronic antigen-induced T cell exhaustion and identifies chromatin remodeling factors as potential targets for improving T cell persistence.

### Mini-pool CRISPR screens validate genetic regulators of T cell exhaustion *in vitro*

To further validate and characterize the top ranked genome-wide screen factors, we created a custom mini-pool of 2,000 sgRNAs, which included sgRNAs that targeted 300 top ranked genes (6 sgRNAs per gene), as well as 100 non-targeting and 100 single-targeting controls (**Table S3**). We repeated the *in vitro* stimulation screen and collected acute and chronic samples, as well as input samples on day 4 (**Table S4**, Figure S6A). We observed high concordance between biological replicates and therefore merged the replicates to perform three comparisons: (1) chronic vs acute, (2) acute vs input, and (3) chronic vs input (Figure S6B-D, **Table S5**). The chronic vs acute comparison served as validation of the original genome-wide screen, and we found that, of the 88 genes in the pool that were positive hits in the genome wide screen, 52 (59.1%) were validated in the mini-pool (FDR < 0.05; Figure 2G, S6D). Next, we compared the chronic vs acute gene enrichments to acute vs input enrichments, which measured the fitness advantage or disadvantage of each gene knockdown in acutely stimulated proliferating T cells in culture (Figure 2H**, left**). Two hits, *Trp53* and *Brd1*, were enriched in both comparisons, demonstrating that depletion of these factors imparts an overall proliferation advantage to T cells in both acute and chronic stimulation conditions. In contrast, most genes displayed either similar (233/300; 77.7%) or reduced (64/300; 21.3%) enrichments in acute stimulation compared to input, enabling the identification of sgRNAs that specifically improve T cell persistence in the presence of chronic antigen, rather than T cell proliferation in general, and that maintain proliferative capacity after acute stimulation (Similar: −3.5 ≤ z ≤ 3, reduced: z < −3.5, improved: z > 3.5; Figure 2H**, left,** S6E). Finally, we compared chronic vs acute sgRNA enrichments to chronic vs input enrichments to identify sgRNAs that have an overall persistence advantage after chronic antigen stimulation, rather than only a comparative advantage to acute stimulation (Figure 2H**, right**). In summary, these mini-pool experiments validated hits from the genome-wide CRISPR screen and identified genes which selectively restrict T cell persistence in the setting of chronic antigen stimulation.

### *In vivo* CRISPR screens identify chromatin remodeling factors that limit T cell persistence in tumors

To characterize the *in vivo* function of the top genome-wide hits, we next screened the sgRNA mini-pool in two murine tumor models. We used Cas9/OT-1 T cells, which recognize the ovalbumin-derived peptide antigen, SIINFEKL, to remove functional T cell variability in the screen due to differing TCR specificities. On day 0, we bilaterally injected MC-38 colon adenocarcinoma or B16 melanoma tumors that ectopically expressed ovalbumin into Rag1^-/-^ mice and isolated CD8^+^ T cells from Cas9/OT-1 mice. On day 1, we transduced the T cells with the custom sgRNA mini-pool (Figure 3A). We transplanted 1×10^6^ T cells per mouse 6 days after tumor inoculation, harvested the tumors and spleens of mice on day 15, sorted T cells from each organ, and sequenced the bulk sgRNA content in these cells (Figure S7A, **Table S4**). We computed sgRNA enrichments as described above and merged the results from all mice to create an aggregate tumor LFC z-score and spleen LFC z-score for each gene in each tumor model, relative to the control distribution (Figure 3B-D, **Table S5**).

**Figure 3:**
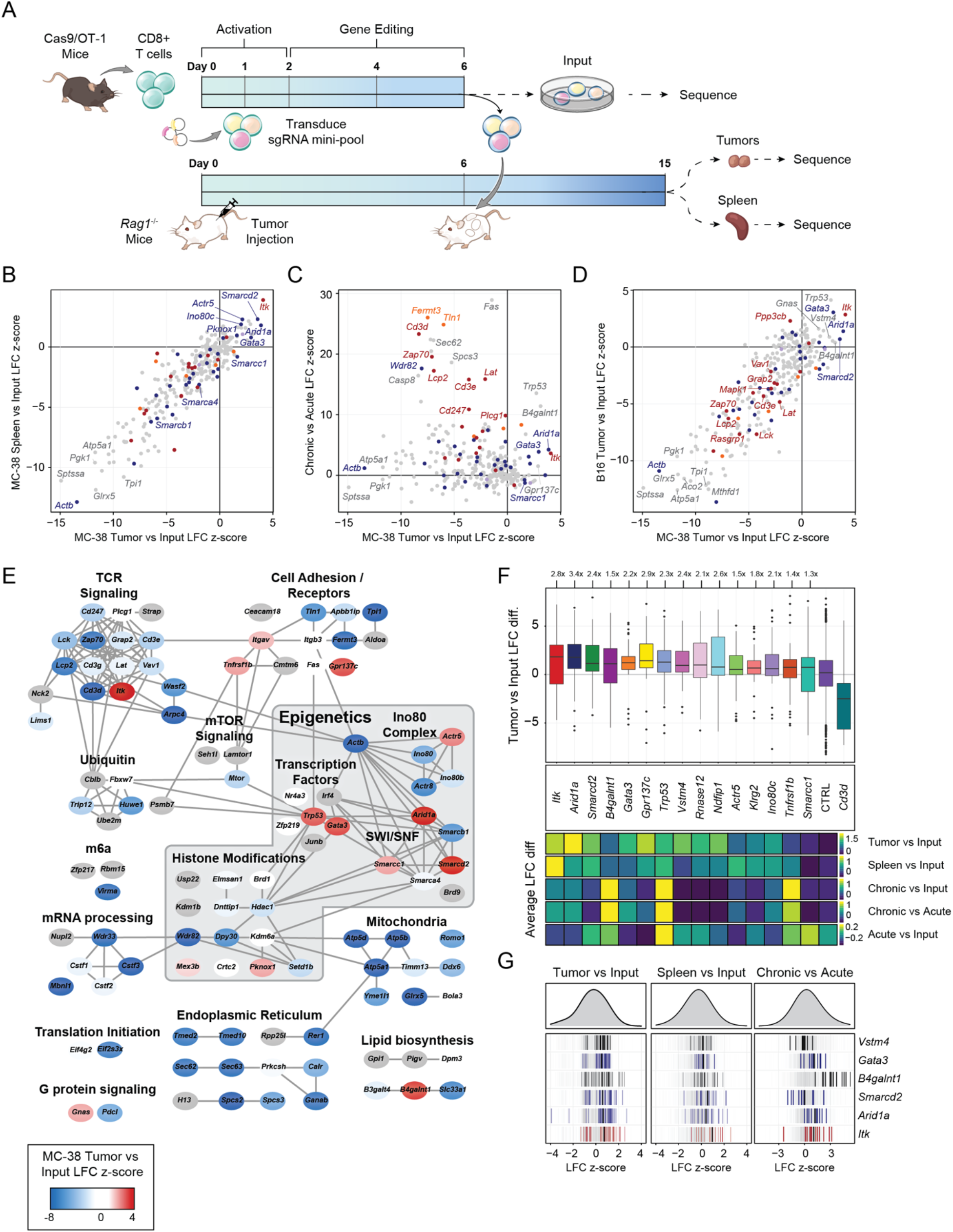
Targeted *in vivo* screening identifies subunits of the INO80 and BAF complexes that limit T cell persistence. **(A)** Diagram of *in vivo* pooled CRISPR screening. **(B)** Correlation of tumor LFC z-scores to spleen LFC z-scores, colored by functional category. **(C)** Correlation of *in vivo* z-score and *in vitro* z-scores for genes in the CRISPR mini-pool. **(D)** Correlation of *in vivo* MC-38 and B16 tumor z-scores for genes in the CRISPR mini-pool. **(E)** Cytoscape protein-protein interaction network colored by z-score in MC-38 tumors. **(F)** Top: Boxplot of tumor versus input log fold change for each sgRNA targeting the indicated gene, with the mean control log fold change subtracted. Bottom: heatmaps showing the sgRNA average of the indicated in vivo or in vitro screen for the same hits. **(G)** Individual sgRNA residuals for six top hits showing the Tumor vs Input comparison (left), Spleen vs Input (center), and *in vitro* Chronic vs Acute (right).

Once again, we first analyzed sgRNAs targeting the TCR complex and signaling genes. In contrast to the *in vitro* screen, cells containing TCR complex/signaling sgRNAs should have an impaired ability to recognize antigen and thus be depleted in tumors. Indeed, sgRNAs targeting nearly all of the previously identified TCR and integrin signaling-related hits were depleted in the tumors and spleens (Figure S7B). Similarly, sgRNAs targeting genes in several other functional categories were also depleted in both tumors and spleens, indicating an impairment of proliferation and/or antigen recognition of these knockdowns *in vivo* (Figure 3B, 3E). Thus, not all categories of genes that improved T cell persistence *in vitro* improved antigen-dependent persistence in the TME *in vivo*, as may be expected due to the multi-factorial process of tumor infiltration and persistence.

However, a select group of *in vitro* hits were highly enriched in tumors and spleens in both tumor models, including *Arid1a*, *Itk*, *Smarcd2*, *B4galnt1*, *Gata3*, *Gpr137c*, *Trp53*, and *Vstm4* (Figure 3B-D). *Gata3* is a transcription factor which was previously demonstrated to regulate the development of T cell exhaustion, and importantly, deletion of this factor improves T cell function, persistence, and tumor control *in vivo* (Singer et al., 2016). The remaining hits, to our knowledge, have not been studied in T cell exhaustion or immunotherapy contexts. Visualizing the tumor enrichments of each gene in the context of the Cytoscape network revealed that many of the positive hits *in vivo* were epigenetic factors, including subunits of two chromatin remodeling complexes, the INO80 complex, (subunits *Ino80c* and *Actr5*), and the BAF complex (subunits *Arid1a, Smarcd2*, and *Smarcc1*; Figure 3E). Other categories were also represented, such as the G-protein coupled receptor, *Gpr137c*, an enzyme involved in ganglioside biosynthesis, *B4galnt1*, and the IL-2 inducible T cell kinase, *Itk*. We calculated the sgRNA enrichments of the top positive hits compared to input controls and found that each gene knockdown improved T cell accumulation in tumors by up to 3.4-fold. For comparison, T cells lacking *Cd3d* were depleted 6.7-fold and T cells lacking *Cd3e* were depleted 3.3-fold, demonstrating that targeting the top hits substantially improved T cell persistence in tumors (Figure 3F, S7C). Furthermore, we observed that the persistence advantage of each knockout was similar in the tumor and the spleen, and no perturbation except *Trp53* displayed substantially improved fitness in the absence of chronic TCR stimulation (during acute stimulation *in vitro*), again demonstrating the specificity of the CRISPR screen strategy to identify perturbations that improve T cell persistence only in the setting of chronic antigen stimulation, rather than improving general T cell fitness (Figure 3F-G, S7C).

### Competition assays and SWI/SNF mini-pool screens demonstrate that tuning cBAF activity can enhance T cell persistence

We sought to validate the persistence advantage of *Arid1a*-sgRNA cells and determine whether these cells retained effector function *in vivo*. We first used a cell competition assay where a single-targeting control (CTRL1) sgRNA was cloned into a retroviral vector expressing a violet-excited fluorescent protein (VEX), while two *Arid1a*-sgRNA sgRNAs (*Arid1a*-1 and *Arid1a-*2) were cloned into a vector which was identical except for the substitution of a blue fluorescent protein (BFP). The activity of both *Arid1a*-targeting sgRNAs was confirmed at the DNA and protein level by Sanger sequencing and Western blot (Figure S8A-C). Cells were separately transduced with either vector, selected with puromycin to enrich for transduced cells, and mixed together. The mixed cells were then put into the *in vitro* chronic stimulation assay (Figure 4A) or the *in vivo* tumor model (Figure 4B). *In vitro* and *in vivo*, *Arid1a*-sgRNA cells demonstrated significantly enhanced persistence, compared to control cells, confirming the results of the pooled screens (Figure 4A-B; *Arid1a*-1 to CTRL1 average normalized ratio: *in vitro* day 10 = 4.03, p = 0.0059; *in vivo* day 15 = 2.46, p = 0.033; *Arid1a*-2 to CTRL1 average normalized ratio: *in vitro* day 10 = 3.79, p = 0.012; *in vivo* day 15 = 2.72, p = 0.0088; Welch Two Sample t-test). Moreover, *Arid1a*-sgRNA cells exhibited lower levels of PD-1 and Tim3 after chronic stimulation *in vitro* (percentage double positive cells: *Arid1a*-1 average decrease of 27.7%, p = 0.00099; *Arid1a*-2 average decrease of 10.6%, p = 0.038; Welch Two Sample t-test; Figure 4A). Finally, we evaluated whether the observed enhanced persistence and altered differentiation trajectory of *Arid1a*-sgRNA cells resulted in improved anti-tumor responses *in vivo*. We inoculated Rag1^-/-^ mice with MC-38 tumors as previously described, and on day 6, transplanted 5×10^5^ Cas9/OT-1 CD8^+^ T cells transduced with either CTRL1 retrovirus or *Arid1a*-sgRNA retrovirus and monitored tumor growth (Figure 4C). By day 15, transfer of *Arid1a*-sgRNA cells significantly improved tumor clearance, compared to transfer of control cells (*Arid1a*-sgRNA vs CTRL1 tumor size, Day 15 p = 5 x 10^-8^; Welch Two Sample t-test). Importantly, survival of mice receiving *Arid1a*-sgRNA T cells was extended by 66%, compared to mice receiving CTRL1 T cells (Median survival = 25 days (*Arid1a*-sgRNA), 12 days (No Transplant), or 15 days (CTRL1); *Arid1a*-sgRNA vs CTRL1 p = 1.20 x 10^-8^; Figure 4D).

**Figure 4:**
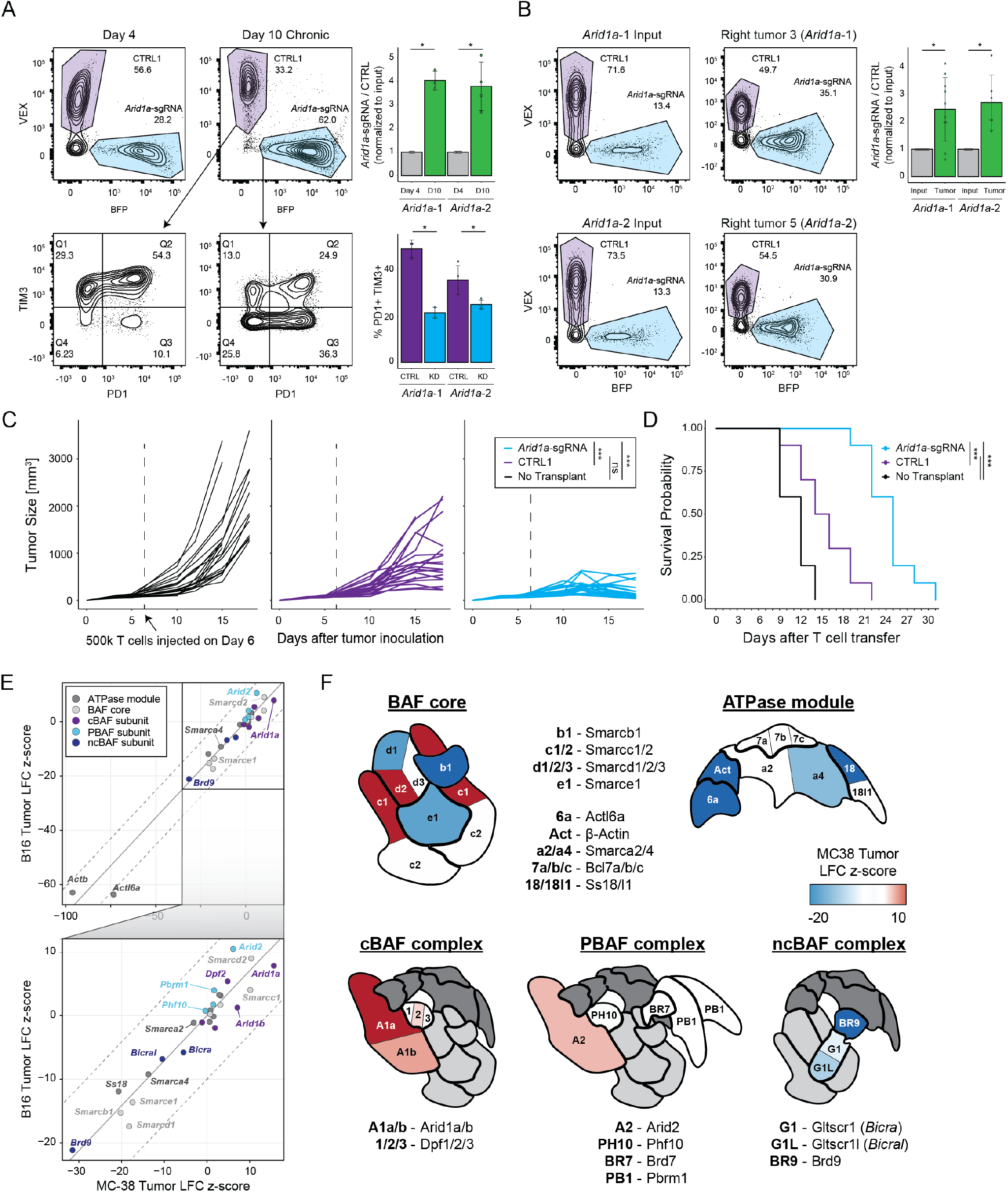
SWI/SNF mini-pool CRISPR screens and functional studies demonstrate that tuning cBAF activity can enhance anti-tumor immunity. **(A)** *In vitro* competition assay of *Arid1a*-sgRNA versus CTRL1 T cells. Left: cells were mixed on Day 4 at the indicated ratios and passaged in the chronic stimulation assay for 6 days. On Day 10, proliferation relative to CTRL1 T cells and surface phenotype were assessed by flow cytometry. **(B)** *In vivo* competition assay of *Arid1a*-sgRNA versus CTRL1 T cells. Cells were mixed on Day 6 (input) and then transplanted into tumor bearing mice. On Day 15, relative proliferation in the tumor was assessed by flow cytometry. **(C)** Tumor sizes for each cohort. Statistical significance was assessed at Day 15. **(D)** *Arid1a*-sgRNA T cells significantly improve survival of tumor-bearing mice. **(E)** Correlation of SWI/SNF CRISPR mini-pool tumor enrichments in MC-38 versus B16 tumor models. **(F)** Cartoons of the three BAF complexes colored by z-score from SWI/SNF CRISPR mini-pool experiments in MC-38 tumors. BAF complex cartoons adapted from (Mashtalir et al., 2018). * p < 0.05, *** p < 0.001.

To provide deeper mechanistic insight into the role of BAF complex factors in T cell exhaustion, we designed an additional CRISPR mini-pool screen targeting each of the 29 SWI/SNF complex subunit genes in the B16 and MC-38 tumor models (described above) and interpreted these results in the structural context of SWI/SNF complex assembly (Mashtalir et al., 2018) (**Table S3**). As observed in the prior *in vivo* screen, the three most significant hits were in the cBAF complex (*Arid1a*, *Smarcc1*, and *Smarcd2*) and notably were in positions of the complex that can be substituted by paralogs in other forms of the complex (Figure 4E-F; **Table S6-7**) (Mashtalir et al., 2018). In contrast, perturbation of irreplaceable subunits of the BAF core (e.g. *Smarce1*, *Smarcb1*) or ATPase module components was deleterious and led to depletion of these sgRNAs. Therefore, we propose a model in which tuning (reducing) the presence of cBAF on chromatin is beneficial for T cell persistence. This concept is supported by prior mechanistic studies demonstrating that *ARID1A*-deficient tumors exhibit reduced (but not ablated) levels of cBAF complex on chromatin, which results in decreased access of key transcription factors (including AP-1 factors) (Mathur et al., 2017; Xu et al., 2020). In addition to cBAF, we also observed positive enrichments of sgRNAs targeting the PBAF complex member, *Arid2*, and strong depletion of sgRNAs targeting the ncBAF complex members, *Bicral, Bicra, and Brd9* (Figure 4E-F; **Table S7**). In summary, these results demonstrate that perturbation of cBAF complex subunit genes, including *Arid1a*, can improve T cell persistence and anti-tumor immunity *in vivo*.

### Perturbation of *ARID1A* improves T cell persistence in primary human T cells

We asked whether perturbation of cBAF subunits could also improve T cell persistence and reduce exhaustion in primary human T cells. We first sought to replicate the *in vitro* chronic stimulation assay using human T cells (Figure 5A). We introduced CRISPR-Cas9/sgRNA RNPs targeting *ARID1A* (two independent sgRNAs) or a control RNP into primary human T cells. We split the cells into acute and chronic cultures, and the chronic condition was stimulated for 6 days with anti-CD3 coated plates (analogous to the mouse assay). In acutely stimulated cultures, we observed no difference between the genotypes for proliferation or viability. However, in chronically stimulated cultures, *ARID1A*-sgRNA cells proliferated significantly more and maintained higher viability than CTRL T cells (*ARID1A*-sgRNA vs CTRL1 cells: mean increase of 22.75% viability, p = 1.70 x 10^-5^, and mean increase of 5.25-fold expansion, p = 0.013; Figure 5A).

**Figure 5:**
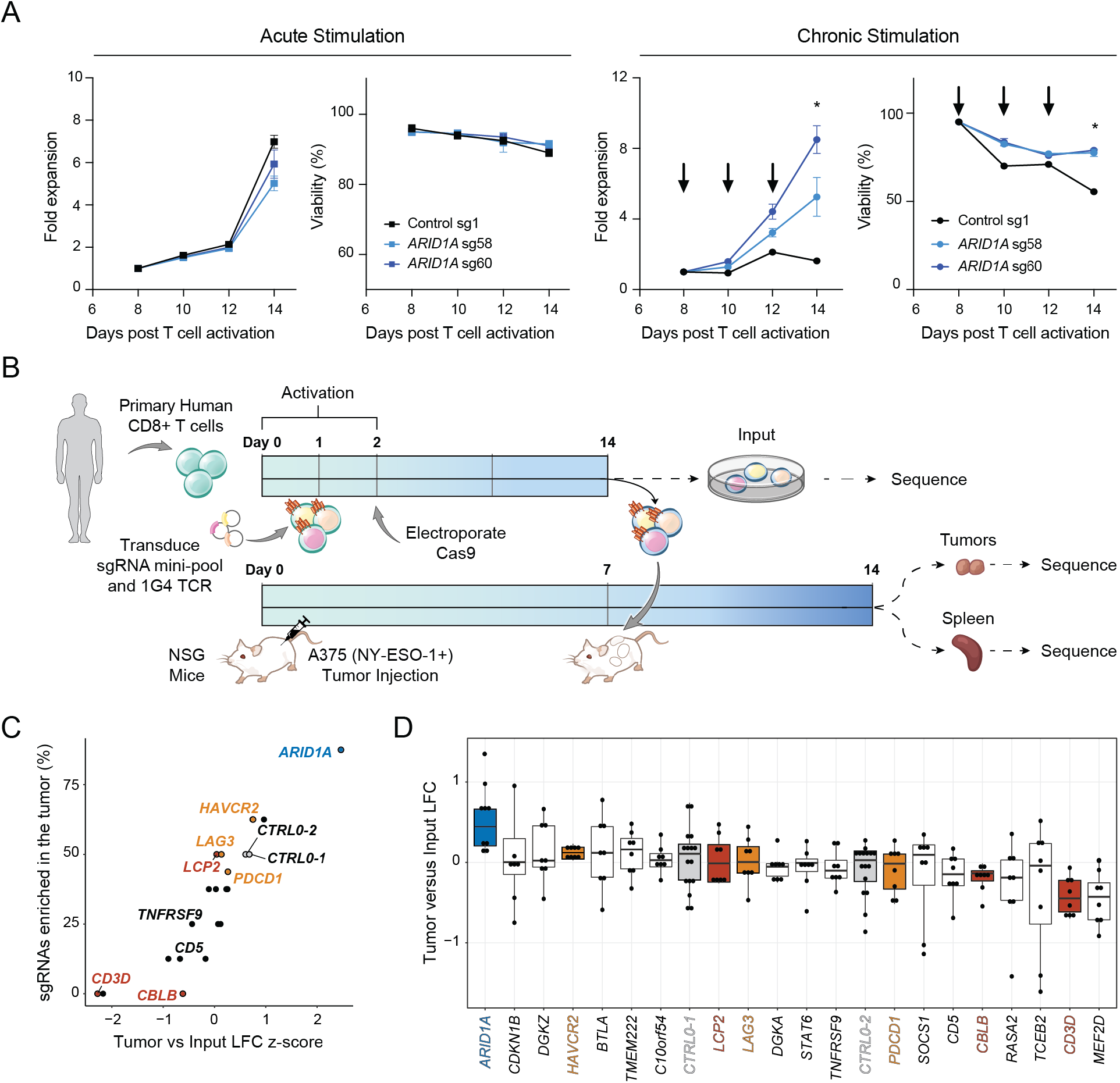
Conserved function of *ARID1A* in human T cells *in vitro* and *in vivo*. **(A)** Proliferation and viability of primary human T cells after electroporation of the indicated RNP. Left: Acutely stimulated T cells. Right: Chronically stimulated T cells using anti-CD3-coated plates. Data shown is representative of 3 independent experiments and 3 donors. **(B)** Schematic of CRISPR mini-pool screen in primary human CD8^+^ T cells transduced with the NY-ESO-1-specific TCR, 1G4. **(C)** Results of the human CRISPR mini-pool screen aggregated by gene. **(D)** Results of the human CRISPR mini-pool screen with individual sgRNAs shown as dots. Genes are ordered from highest to lowest average LFC. Results shown are merged from 2 independent donors and 2 mice per donor.

We next sought to validate the persistence advantage of *ARID1A*-sgRNA T cells *in vivo* and in the context of other genetic factors that have recently emerged from human T cell functional CRISPR screens (Shifrut et al., 2018). We designed a CRISPR mini-pool for *in vivo* human T cell experiments, which encompassed 48 sgRNAs targeting 20 genes and included 16 negative control guides (**Table S8**). We included sgRNAs targeting *ARID1A*, as well as the inhibitory receptors, *PDCD1*, *LAG3*, and *HAVCR2*, and other top-ranked genes from our prior screens, such as *TMEM222, CBLB, TCEB2,* and *SOCS1* (Shifrut et al., 2018). We performed the screen in the A375 human melanoma xenograft model, which expresses the NY-ESO-1 antigen that can be targeted with the 1G4 TCR. We introduced the cognate 1G4 TCR into primary human T cells from two independent donors on day 1 along with the sgRNAs, and on day 14 transplanted T cells into NOD-SCID-IL2Rγ-null (NSG) tumor-bearing mice (Figure 5B). 7 days later, we sorted T cells from the tumors and spleens, sequenced sgRNAs present in each organ, and compared their abundance to input samples prior to transplant. As expected, we did not observe enrichments in control sgRNAs or sgRNAs targeting inhibitory receptors, while we did observe depletion of sgRNAs targeting *CD3D* (Figure 5C-D). In contrast, and consistent with our results in murine T cells, sgRNAs targeting *ARID1A* were significantly enriched in tumors compared to input samples in both donors, demonstrating that the function of cBAF in limiting T cell persistence is also conserved in human T cells (7 of 8 *ARID1A*-sgRNA sgRNAs enriched in tumors versus input; *ARID1A*-sgRNA versus CTRL LFC p = 0.0010 by Wilcoxon test; Figure 5C-D).

### *In vivo* Perturb-seq reveals distinct transcriptional effects of chromatin remodeling complexes in TILs

To understand the molecular mechanisms driving improved T cell function in hits identified by the *in vitro* and *in vivo* CRISPR screens, we performed Perturb-seq, which simultaneously captures CRISPR sgRNAs and the transcriptome in single cells (Adamson et al., 2016; Dixit et al., 2016; Replogle et al., 2020). We designed a third custom sgRNA pool (micro-pool) targeting the INO80 and BAF complexes. Both complexes are ATP-dependent chromatin remodelers that are essential in many aspects of development (Hargreaves and Crabtree, 2011). For SWI/SNF genes, we targeted *Arid1a*, *Smarcc1*, and *Smarcd2* (top hits identified *in vitro* and *in vivo)*, as well as *Arid2* and *Arid1b*, which were enriched in the SWI/SNF-specific mini-pool screen. Of these, *Smarcc1* and *Smarcd2* are in the BAF core, *Arid1a* and *Arid1b* are in the cBAF complex, and *Arid2* is present only in the PBAF complex. From the INO80 complex, we selected *Actr5* and *Ino80c*, which were enriched in both the *in vitro* and *in vivo* screens. Interestingly, the yeast homologues of *Actr5* and *Ino80c*, Arp5 and Ies6, have been shown to physically associate with each other, forming a subcomplex independent of the rest of the INO80 complex. The subcomplex can modulate the activity of the rest of the INO80 complex; it interacts with chromatin in an INO80-dependent manner and repositions nucleosomes (particularly the +1 nucleosome) to activate gene transcription, especially at metabolism-related genes (Yao et al., 2016). Finally, we included positive controls, *Pdcd1* and *Gata3*, as well as 12 single targeting negative controls for a total of 48 sgRNAs targeting 9 genes (**Table S3**). We performed a similar *in vivo* T cell protocol as described above for the larger CRISPR screen: we isolated CD8^+^ T cells from Cas9/OT-1 mice, transduced these cells with the sgRNA micro-pool, and then transplanted them into Rag1^-/-^ mice bearing MC-38 ovalbumin tumors. As in the prior screens, we also collected an input sample (collected on the day of transplantation) to evaluate the persistence phenotype of each sgRNA. Nine days after T cell transplantation, we harvested tumors, isolated TILs, and used direct-capture Perturb-seq to read out sgRNA identity and gene expression profiles simultaneously using the 10x Genomics 5’ gene expression platform (Figure 6A) (Replogle et al., 2020). We sequenced cells from seven biological replicate Perturb-seq samples across two independent experiments (Figure S9A-B).

**Figure 6:**
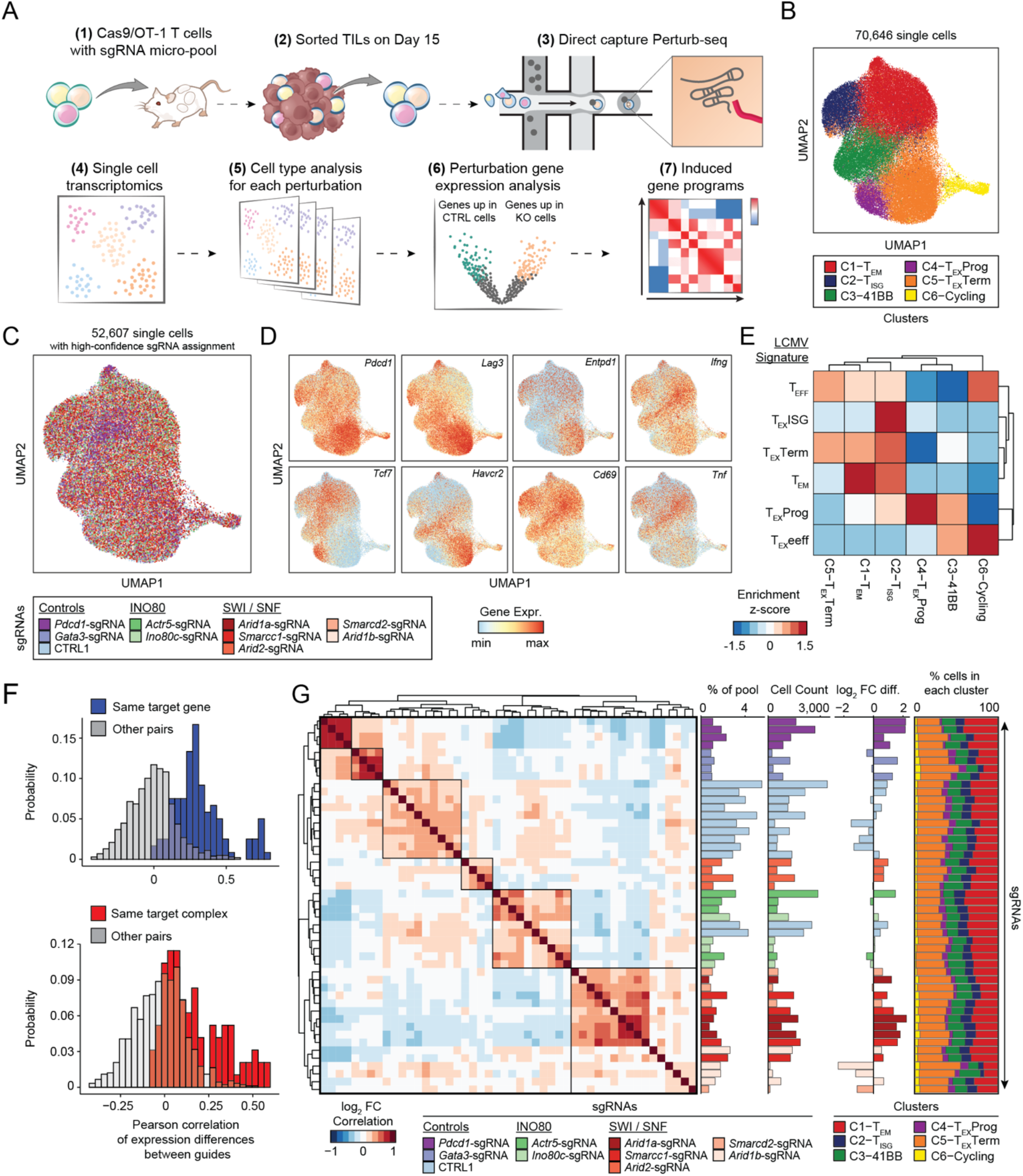
*In vivo* Perturb-seq reveals distinct transcriptional roles of the cBAF and INO80 complexes in TILs. **(A)** Diagram of direct-capture Perturb-seq of sorted TILs. **(B)** scRNA-seq profiles of TILs colored by cluster assignment. **(C)** scRNA-seq profiles of cells colored by the perturbation detected in each cell. Cells where no guide, or multiple guides, were detected are shown in grey. **(D)** Expression of selected marker genes in each single cell. **(E)** Analysis of LCMV signature gene sets for each cluster. Gene set enrichment scores were calculated for each single cell, cell values were averaged by cluster and z-scored. **(F)** Histogram of Pearson correlation of gene expression differences of pairs of sgRNAs. Top: Pairs targeting the same gene are shown in blue, other pairs are shown in gray. Bottom: Pairs targeting the same protein complex are shown in red, other pairs are shown in gray. Complexes considered in the analysis are cBAF (*Arid1a*, *Arid1b*, *Smarcd2*, and *Smarcc1*) and INO80 (*Ino80c* and *Actr5*). Pairs of sgRNAs that target the same gene are excluded. **(G)** Left: Heatmap of the correlation of gene expression differences of each pair of sgRNAs. Center (from left to right): Representation of each sgRNA in the pre-transplant sample, cell count of each sgRNA in the Perturb-seq dataset, and estimated fold change of each sgRNA relative to controls. Right: Proportion of cells in each cluster for each sgRNA.

After quality control filtering, we obtained high-quality scRNA-seq profiles from 70,646 cells, and scRNA-seq clustering and dimensionality reduction identified 6 clusters (Figure 6B). We determined a high-confidence sgRNA identity for each cell by using z-scores to quantify the enrichment of each sgRNA, relative to other sgRNAs detected in the same cell. Cells were assigned to a particular sgRNA if that sgRNA had a z-score of at least 5, and a z-score at least 2 units higher than the next-most prevalent sgRNA. With this strategy, cells with multiple enriched sgRNAs due to retroviral infection doublets, single-cell capture doublets, and/or background reads were removed from further analysis, and 52,607 cells were confidently assigned to a single sgRNA (74.4% of the 70,646 total cells across the two independent experiments; Figure 6C). Cell type clusters expressed varied levels of inhibitory receptors, effector cytokines, and key transcription factors, indicating that they represented a mix of exhausted and effector T cells in the TME (Figure 6D, S9C-D). Cluster 1 cells expressed high levels of *Klf2* and *S1pr1* (T effector memory; T_EM_), Cluster 2 expressed high levels of interferon stimulated genes (ISGs) including *Mx1* (T_ISG_), Cluster 3 expressed high levels of *Tnfrsf9* (encoding 41BB) and *Cd160* (T-41BB), Cluster 4 expressed high levels of progenitor exhaustion genes including *Pdcd1*, *Tcf7* and *Slamf6* (T_EX_Prog), Cluster 5 expressed the highest levels of inhibitory receptors *Pdcd1*, *Lag3*, and *Havcr2* (T_EX_Term), and Cluster 6 consisted primary of cycling cells, marked by *Mki67* and confirmed by cell cycle analysis (T-Cycling; Figure S9C-D). To further refine the cluster identities, we generated gene signatures from previously published CD8^+^ T cell types present in acute or chronic LCMV infection *in vivo* (Figure S9E-F) (Daniel et al., 2021). We used the top 100 marker genes for each LCMV T cell cluster to score each single cell in our Perturb-seq dataset according to the average expression of these signature gene sets. Visualizing the enrichment of these LCMV signatures in each cluster demonstrated transcriptional similarity of several clusters to cell types in the reference dataset (Figure 6E). For example, Cluster 1 was enriched for the effector memory-related genes (T_EM_ signature), Cluster 2 was similar to the T_EX_ISG signature, and the progenitor and terminally exhausted clusters (Clusters 4 and 5) enriched the corresponding LCMV signatures (Figure 6E).

We next performed several sgRNA-level quality controls to assess the reproducibility of effects of independent sgRNAs (Figure 6F-G). We computed gene expression differences between each sgRNA and all other cells in the dataset and confirmed that independent sgRNAs targeting the same gene had highly correlated gene expression changes relative to pairs of sgRNAs targeting different genes, which, as expected, centered around zero (Figure 6F top). We next asked whether this similarity also extended to sgRNAs targeted distinct genes in the same protein complex. We evaluated the correlation of pairs of sgRNAs targeting the same complex, grouping together guides targeting cBAF genes (*Arid1a*, *Arid1b*, *Smarcc1*, and *Smarcd2*) and guides targeting INO80 genes (*Ino80c* or *Actr5*). Strikingly, we found that these sgRNA pairs were also significantly more correlated than all pairs of guides, indicating common transcriptional effects of targeting distinct subunits within the same complex (Figure 6F **bottom**). We next visualized gene expression correlations of all pairs of sgRNAs together (Figure 6G). Unbiased clustering organized sgRNAs into correlated groups, primarily driven by target gene and target complex identity. Interestingly, *Arid2* clustered separately from the rest of the BAF targeted sgRNAs, suggesting distinct roles for the cBAF and PBAF complexes (Figure 6G). We next used the input representation of each sgRNA along with the number of cells detected with each sgRNA to estimate the T cell accumulation advantage between each sgRNA and the set of single targeting negative controls (Figure 6G). This analysis demonstrated that the majority of sgRNAs enhanced T cell accumulation in the tumor, relative to control sgRNAs, in line with the *in vivo* screen results. In particular, *Arid1a*-sgRNA cells were enriched 2.74-fold on average relative to controls, while *Pdcd1*-sgRNA cells were enriched 2.67-fold on average relative to controls (Figure 6G). Finally, we examined the cell type cluster composition of cells containing each sgRNA (Figure 6G**, far right**). Notably, all perturbations contained cells from each cluster with relatively similar proportions, suggesting that depletion of each target gene may not impact wholescale changes in cell type compositions or trajectories, but rather modulate gene expression in one or more clusters.

To further investigate this possibility, we aggregated cells that contained sgRNAs targeting the same gene and computed differential genes for each perturbation, compared to CTRL1 cells (Figure 7A). Targeting cBAF subunits *Arid1a*, *Smarcd2*, or *Smarcc1* induced shared global changes in the transcriptional program of T cells, including the upregulation of effector molecules, *Gzmb* and *Ifng*, cell surface receptors, *Cxcr6* and *Il7r*, and transcription factors, *Irf4* and *Batf*. Meanwhile, *Pdcd1*, *Lag3*, and *Ccl5* were consistently downregulated by cBAF perturbation (Figure 7A). In contrast, *Arid2* perturbation induced a distinct gene expression program, albeit with some similarities, including the downregulation of *Pdcd1* and *Lag3*. Perturbation of *Gata3* and *Pdcd1* induced distinct gene expression changes from either cBAF or *Arid2* perturbation; for example, the most upregulated gene after *Pdcd1* depletion was *Tox*, consistent with the proposed impact of PD-1 deletion on accelerating differentiation to terminal exhaustion (Figure 7A). To quantify the aggregate similarity of gene expression changes induced by each perturbation, we correlated all pairs of perturbations and performed clustering to group perturbations that were similar according to this metric (Figure 7B). This analysis quantitatively confirmed our observation that cBAF perturbations, *Arid1a*, *Smarcc1*, and *Smarcd2*, induced similar programs, while INO80 perturbations, *Ino80c* and *Actr5,* also exhibited highly correlated changes (that are distinct from that induced by cBAF perturbation). In contrast, *Pdcd1* and *Gata3* perturbations clustered separately, although with moderate correlation to each other. Finally, when gene expression changes were considered for perturbed vs CTRL1 cells within each cluster, we found that each perturbation induces highly concordant changes in gene expression regardless of the T cell subtype (Figure S9G).

**Figure 7:**
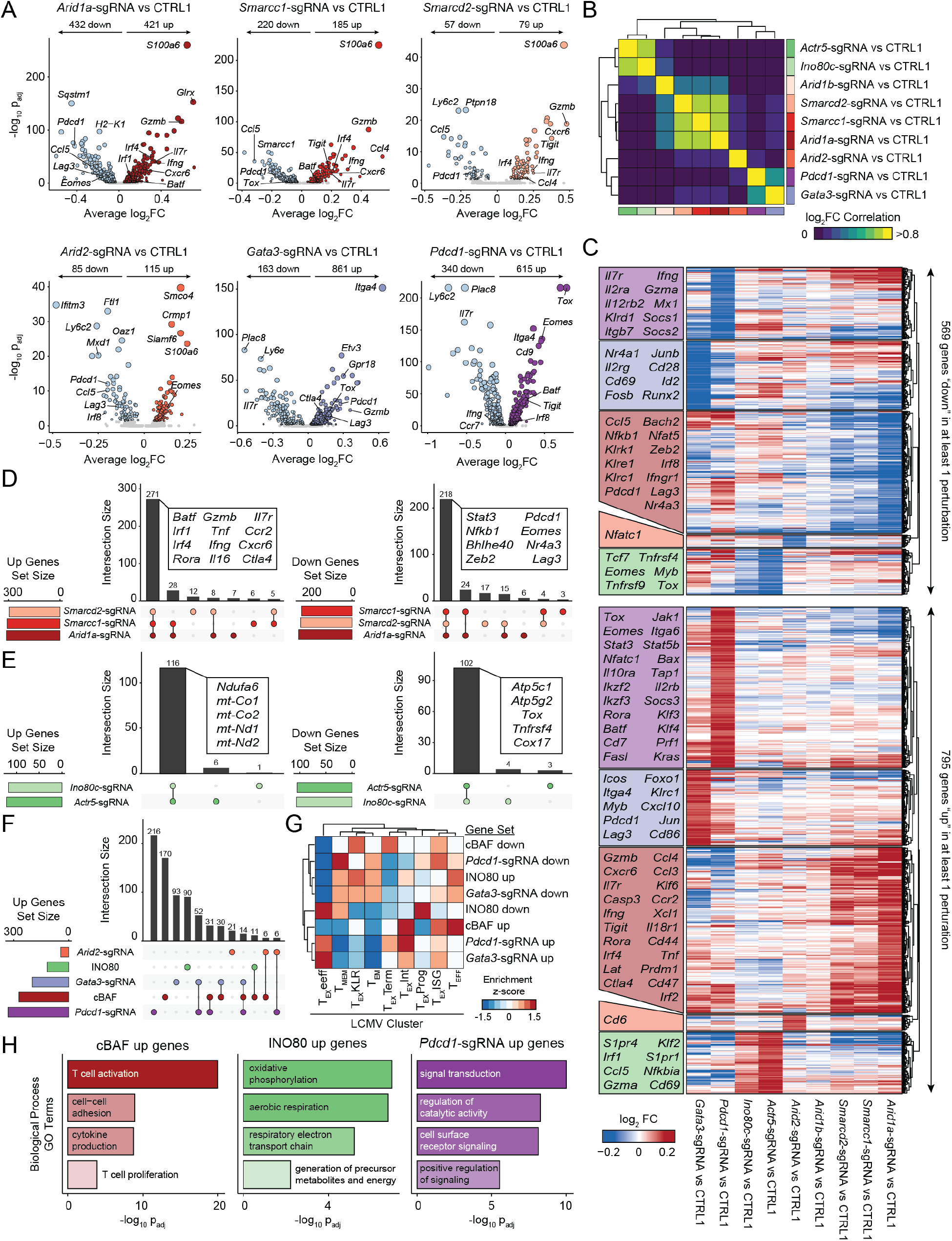
cBAF-depleted T cells exhibit enhanced effector gene signatures and reduced terminal exhaustion. **(A)** Volcano plots comparing aggregated cells with the indicated perturbation versus CTRL1 cells. **(B)** Pairwise correlations of gene expression differences induced by each perturbation. **(C)** Heatmap of all upregulated (up) or downregulated (down) genes in at least one perturbation, grouped by which perturbation has the strongest effect on expression. Selected genes in each block are labeled. **(D)** Comparison of upregulated or downregulated gene sets by perturbation of cBAF subunits, *Arid1a*, *Smarcd2*, or *Smarcc1*. **(E)** Comparison of gene sets up- or downregulated by perturbation of INO80 subunits *Actr5*, or *Ino80c*. **(F)** Comparison of gene sets upregulated by perturbation of cBAF subunits, INO80 subunits, or *Pdcd1*, *Gata3*, or *Arid2*. **(G)** Enrichments of upregulated and downregulated gene sets in LCMV expression data (Daniel et al., 2021). Module scores of each gene set were computed for each single cell in the LCMV dataset, averaged by cluster, and then z-scored to obtain the indicated enrichment z-scores. **(H)** Selected GO Terms of indicated gene sets.

We aggregated all genes significantly differential between perturbed cells and CTRL1 cells and defined core gene programs perturbed by depletion of the cBAF and INO80 complexes (Figure 7C-E). We found that the upregulated and downregulated gene sets were highly conserved within each complex (Figure 7D-F, S10A), with cBAF perturbation inducing genes such as *Batf*, *Irf4*, *Il7r*, and *Ccr2*, while repressing genes such as *Stat3*, *Nfkb1*, *Nr4a3,* and *Eomes*. In contrast, INO80 perturbation substantially modulated metabolism related genes (Figure 7E). Projection of genes upregulated by cBAF depletion onto canonical T cell states identified in chronic LCMV infection showed an enrichment in effector T cell genes, while projection of downregulated genes showed an enrichment in terminal exhaustion-related genes (Figure 7G, S10B). Finally, we performed GO Term analysis of upregulated gene sets. Genes upregulated in cBAF-deficient T cells enriched terms, including T cell activation, cell adhesion, cytokine production, and T cell proliferation, while genes upregulated in INO80-deficient T cells enriched metabolic terms, including oxidative phosphorylation and aerobic respiration (Figure 7H). In contrast, perturbation of *Pdcd1* induced cell signaling related terms (Figure 7H). In summary, these data demonstrate that these subunits of the cBAF and INO80 chromatin remodeling complexes have distinct roles in T cell exhaustion that are largely conserved within the same complex, with cBAF primarily regulating effector- and exhaustion-related genes and INO80 regulating metabolism. Furthermore, the transcriptional impact of targeting chromatin remodeling factors are minimally overlapping with the impact of previously known targets, *Pdcd1* and *Gata3*, suggesting the potential to synergistically target multiple factors to improve T cell function (Figure 7F, S10A).

### *Arid1a* perturbation limits the acquisition of terminal exhaustion-associated chromatin accessibility

We next asked how perturbation of the cBAF complex via targeting *Arid1a* impacted the epigenetic landscape of T cell exhaustion. We performed a competition assay as described above, wherein CTRL1 and *Arid1a*-sgRNA cells were mixed at a defined ratio and subjected to *in vitro* exhaustion. We used two independent sgRNAs targeting *Arid1a* in duplicates for a total of four replicate samples. At Day 6 and Day 10, we isolated CTRL1 and *Arid1a*-sgRNA cells from the same culture and performed ATAC-seq on each population. To analyze these results in the context of our initial assay characterization (Figure 1), we included the profiles of naïve (Day 0) and activated (Day 2) WT T cells (Figure 8A). The chromatin state progression in CTRL1 cells proceeded similarly to that observed previously in unperturbed cells; however, *Arid1a*-sgRNA cells proceeded down a distinct chromatin state trajectory, remaining closer to naïve and activated samples than the CTRL1 cells at both time points (Figure 8A).

**Figure 8:**
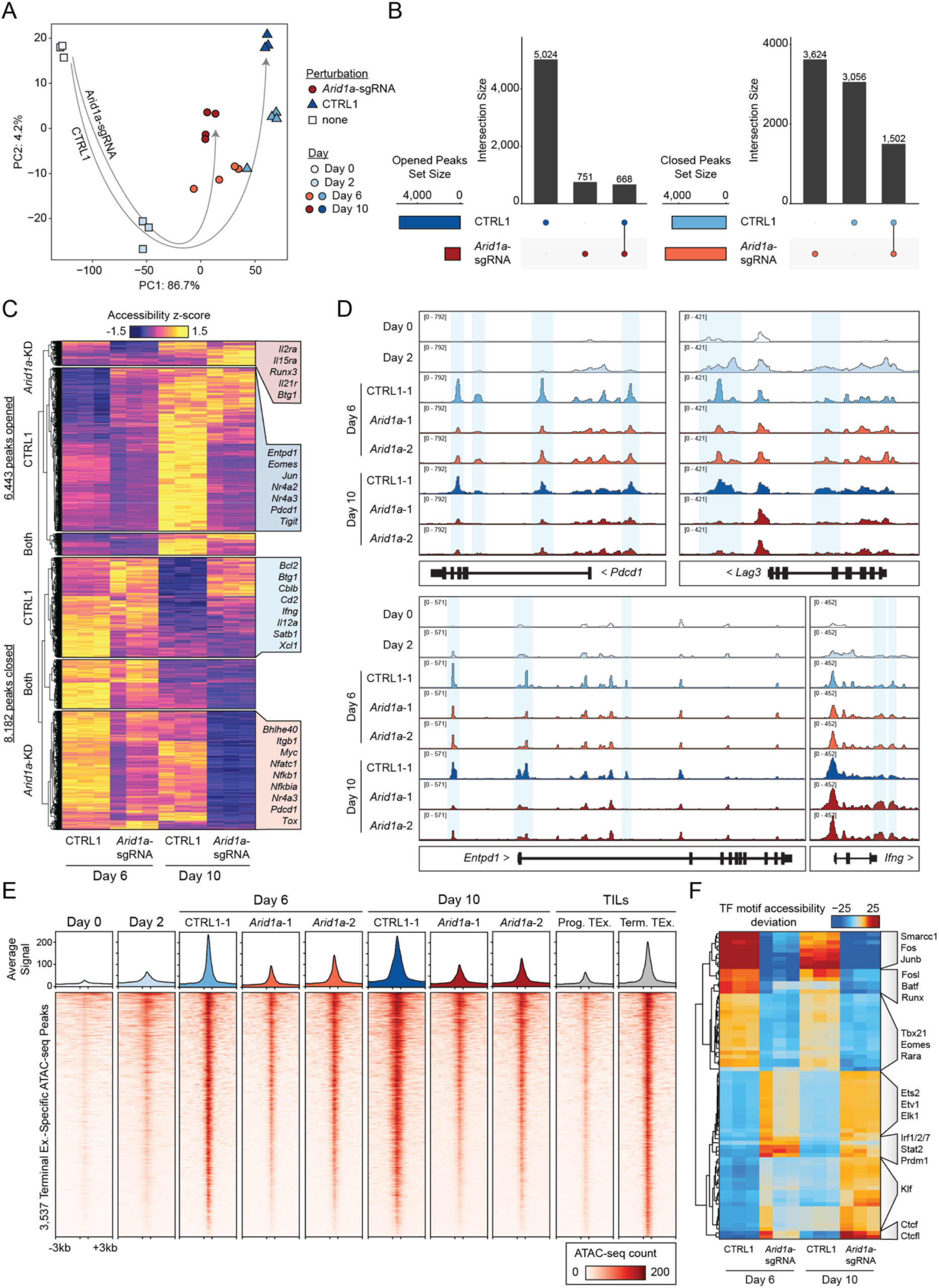
*Arid1a* is required for the acquisition of the exhausted T cell chromatin state. **(A)** Principal component analysis of ATAC-seq profiles of *Arid1a*-sgRNA and CTRL1 cells in the *in vitro* exhaustion competition assay. Unperturbed naïve and activated samples (Day 0 and 2) are included for reference. **(B)** Comparison of ‘opened’ and ‘closed’ ATAC-seq peak sets from Day 6 to Day 10 for each genotype. **(C)** Visualization of ‘opened’ and ‘closed’ ATAC-seq peak sets, with selected nearest genes labeled. **(D)** ATAC-seq signal tracks of selected gene loci. Representative replicates are shown for each condition. **(E)** Heatmap showing ATAC-seq coverage of each peak in the “Terminal Exhaustion peak set” for *Arid1a*-sgRNA and CTRL1 cells at Day 6 and Day 10 in the *in vitro* exhaustion assay. Reference data from TILs is also included, as well as reference naïve and activated cell profiles. **(F)** chromVAR motif accessibility heatmap for *Arid1a*-sgRNA and CTRL1 ATAC-seq samples. Selected motifs are indicated on the right. Top 100 most variable motifs are shown.

Analysis of dynamic chromatin accessibility changes between Day 6 and 10 revealed striking differences between *Arid1a*-sgRNA and CTRL1 cells. We categorized regulatory elements as ‘opened’ peaks, which represented those with increased accessibility at Day 10, compared to Day 6, while ‘closed’ peaks represented elements with decreased accessibility at Day 10, compared to Day 6 (p_adj_ < 0.05, Log_2_ FC > 1). *Arid1a*-sgRNA and CTRL1 T cells demonstrated substantially different chromatin remodeling by comparing these peak sets (Figure 8B-C). First, *Arid1a*-sgRNA cells exhibited a marked decrease in the number of opened peaks, likely representing an impairment of cBAF-depleted cells for opening new regulatory elements (Opened peaks: 1,419 (*Arid1a*-sgRNA) versus 5,692 (CTRL1); Figure 8B). Second, while *Arid1a*-sgRNA cells and CTRL1 cells closed chromatin to a similar extent, the majority of these regions were non-overlapping (Closed peaks: 5,126 (*Arid1a*-sgRNA) versus 4,558 (CTRL1); Figure 8B). Inspecting individual gene loci revealed changes in accessibility in the *Pdcd1*, *Lag3*, *Entpd1*, and *Ifng* loci, demonstrating several open chromatin sites in CTRL1 cells with substantial loss of accessibility in *Arid1a*-sgRNA cells (Figure 8D). Analyzing the terminal exhaustion-specific regulatory element set defined in Figure 1, we found that these sites were less accessible in *Arid1a*-sgRNA cells than in CTRL1 cells at Day 6 and Day 10 (Figure 8E). Finally, we analyzed chromatin accessibility at TF binding sites using chromVAR, which showed that terminal exhaustion-associated TF motifs, including Fos, Jun, and Batf motifs were significantly less accessible in *Arid1a*-sgRNA cells, compared to CTRL1 cells (Figure 8F). Conversely, several TF motifs associated with effector T cell function, including Ets, Klf, and Irf motifs, showed increased accessibility in *Arid1a*-sgRNA cells. In summary, these results suggest that depletion of cBAF and *Arid1a* may improve T cell function by preventing the acquisition of the terminal exhaustion-associated chromatin state.

## Discussion

In this study, we performed genome-wide CRISPR screens in chronically stimulated T cells, which provide a comprehensive atlas of genes that regulate T cell exhaustion. We used a complementary *in vitro* and *in vivo* screening strategy: (1) the development of an *in vitro* exhaustion assay that is compatible with genome-wide CRISPR screening enabled us to massively scale the number of cells, and therefore sgRNA library coverage, compared to prior screens, providing an unbiased and systematic discovery tool, and (2) *in vivo* follow-up screens identified perturbations that significantly improved T cell persistence in immunotherapy-relevant tumor models. Importantly, this strategy recovered known regulators of exhaustion, including *Gata3*, which has previously been demonstrated to limit T cell function in mouse and human tumor models (Singer et al., 2016). However, these screens also uncovered new genes, with a surprising enrichment of epigenetic factors involved in chromatin and nucleosome remodeling, including the cBAF and INO80 complexes, which limited T cell persistence in many cases to a degree greater than previously identified genes. *In vivo* Perturb-seq experiments revealed that depletion of cBAF and INO80 complex subunits impacted distinct gene programs; cBAF modulation led to the upregulation of an effector program and downregulation of terminal exhaustion genes, while INO80 modulation primarily impacted gene expression related to metabolic function. Finally, depletion of the cBAF complex subunit *Arid1a* improved T cell persistence in *in vitro* and *in vivo* competition assays and improved anti-tumor immunity after adoptive T cell transfer.

Our strategy was to isolate a key determinant of T cell dysfunction in cancer — chronic stimulation through the TCR — from the multifactorial process involving tumor localization, trafficking, and immunosuppressive effects in the TME, which may occur to varying degrees across cancer types. The advantage of this strategy is its specificity; sgRNA abundance is impacted only by a single selection factor, and therefore, we provide a precise conceptual picture of the molecular drivers of T cell exhaustion. For example, T cell inhibitory receptors such as PD-1 and CTLA-4 were not hits in our screen, supporting the growing notion that checkpoint blockade does not work by reversing or preventing the process of T cell exhaustion, but rather by recruiting new functional T cell clones to enter the TME (Yost et al., 2019; Pauken et al., 2016; Spitzer et al., 2017; Yost et al., 2021). However, we wish to acknowledge that this strategy does not account for additional dysfunction pathways in T cells that may be mediated by other external stimuli, for example TGFβ-mediated suppression, or specific metabolic or nutrient stressors (Mariathasan et al., 2018; DePeaux and Delgoffe, 2021). Similarly, our follow-up *in vivo* screen selected for one functional aspect of exhaustion — T cell persistence in tumors — but did not account for additional aspects, such as cytokine secretion, and thus, additional genes identified in the *in vitro* screen may be uncovered as important regulators of other facets of T cell exhaustion in future studies.

The enrichment of chromatin remodeling factors as hits in both *in vitro* and *in vivo* screens provides a key complementary message to previous epigenomic profiling studies on T cell exhaustion (Pauken et al., 2016; Sen et al., 2016; Philip et al., 2017; Scott-Browne et al., 2016). Namely, these prior studies demonstrated that exhaustion is mediated by global chromatin remodeling, which maintains a stable dysfunctional cellular phenotype that is not changed by anti-PD-1 treatment. We now demonstrate targeting nucleosome remodeling complexes may be sufficient to prevent the acquisition of features of this exhaustion-associated chromatin state, and thereby improve T cell persistence and maintenance of an effector-like state. It is possible that deletion of these factors may function to ‘dampen’ the downstream epigenetic impact of chronic TCR signaling and thereby extend the window in which T cells can engage antigens without accumulating deleterious terminal exhaustion-associated epigenetic changes. Recent studies in fibroblasts have demonstrated that AP-1 family TFs may have an essential role in signal-dependent enhancer selection by collaboratively binding to nucleosomal enhancers and recruiting the BAF complex to establish accessible chromatin (Vierbuchen et al., 2017). Indeed, analysis of *Arid1a*-sgRNA T cells revealed a dramatic loss of accessibility at exhaustion-induced AP-1 motif-containing regulatory elements, suggesting a similar mechanism in T cells. Importantly, in recent years, the identification of key molecular drivers of T cell exhaustion have been rapidly translated to develop improved clinical-grade cellular immunotherapies, particularly in the context of solid tumors, where exhaustion is a major barrier to therapeutic efficacy (Chen et al., 2019; Lynn et al., 2019; Weber et al., 2021). We envision that future work will build upon these findings to pursue novel avenues and targets to effectively improve T cell function in the context of cancer immunotherapy.

## Acknowledgement

We thank Dhananjay Wagh and the Stanford Functional Genomics Facility for assistance with sequencing. We thank Jasmine Sosa, Ian Lai, and the Stanford Transgenic, Knockout, and Tumor model Center (TKTC) for assistance with tumor experiments. This work was supported by the National Institutes of Health grants U01CA260852 (A.T.S.) and UM1HG012076 (A.T.S.), the Burroughs Wellcome Fund Career Award for Medical Scientists (A.T.S. and S.A.V.), the Parker Institute for Cancer Immunotherapy (A.T.S.), a Pew-Stewart Scholars for Cancer Research Award (A.T.S.), a Cancer Research Institute Technology Impact Award (A.T.S.), a Baxter Foundation Faculty Scholar Award, and the Stanford Innovative Medicine Accelerator and Stanford ChEM-H (A.T.S). A.M. has received the Burroughs Wellcome Fund Career Award for Medical Scientists and The Cancer Research Institute (CRI) Lloyd J. Old STAR award. The Marson lab has received support from the Parker Institute for Cancer Immunotherapy (PICI), the Innovative Genomics Institute (IGI), and the Chan Zuckerberg Biohub. J.A.B was supported by a Stanford Graduate Fellowship and the National Science Foundation Graduate Research Fellowship under Grant No. DGE-1656518. S.A.V. was supported by a Special Fellow Award from the Parker Institute for Cancer Immunotherapy and a Mentored Clinical Scientist Career Development Award from NCI/NIH (K08 CA237731).

## Author contributions

J.A.B. and A.T.S. conceived the study. J.A.B., W.Y., N.L., Y.C., Q.S., A.M.V., C.V.D., M.A.H., Z.M., and S.A.V. performed experiments. K.A.F., E.S., N.K., and J.C. performed experiments in primary human T cells. A.A., C.M., A.M., and J.C. provided resources and supervision for experiments with primary human T cells. J.A.B. analyzed data. K.E.Y. assisted with LCMV data. J.A.B. and A.T.S. supervised all experiments and wrote the manuscript. All authors reviewed and provided comments on the manuscript.

## Declaration of interest

A.T.S. is a scientific co-founder of Immunai and founder of Cartography Biosciences and receives research funding from Arsenal Biosciences, Allogene Therapeutics, and Merck Research Laboratories. J.A.B. is a consultant to Immunai. S.A.V. is an advisor to Immunai. K.E.Y. is a consultant to Cartography Biosciences. C.L.M. is a co-founder of Lyell Immunopharma and Syncopation Life Sciences, and consults for Lyell, Syncopation, NeoImmune Tech, Apricity, Nektar, Immatics, Mammoth and Ensoma. A.A. is a co-founder of Tango Therapeutics, Azkarra Therapeutics, Ovibio Corporation, and Kytarro; a consultant for SPARC, Bluestar, ProLynx, Earli, Cura, GenVivo, Ambagon, Phoenix Molecular Designs and GSK; a member of the SAB of Genentech, GLAdiator, Circle and Cambridge Science Corporation; receives research support from SPARC and AstraZeneca; holds patents on the use of PARP inhibitors held jointly with AstraZeneca. A.M. is a co-founder of Spotlight Therapeutics, Arsenal Biosciences, and Survey Genomics. A.M. is a member of the scientific advisory board of NewLimit. A.M. owns stock in Arsenal Biosciences, Spotlight Therapeutics, NewLimit, Survey Genomics, PACT Pharma, and Merck. A.M. has received fees from 23andMe, PACT Pharma, Juno Therapeutics, Trizell, Vertex, Merck, Amgen, Genentech, AlphaSights, Rupert Case Management, Bernstein, and ALDA. A.M. is an investor in and informal advisor to Offline Ventures and a client of EPIQ. The Marson lab has received research support from Juno Therapeutics, Epinomics, Sanofi, GlaxoSmithKline, Gilead, and Anthem. K.A.F., E.S., J.C., A.A., A.M., and C.L.M. hold patents in the arena of CAR T cell therapeutics. J.A.B. and A.T.S. have filed a patent related to the contents of this study.

### Data and materials availability

CRISPR screen counts tables and z-score tables are available with the manuscript as supplemental data. ATAC-seq and scRNA-seq data will be deposited on GEO.

**Table S1:** sgRNA counts for genome-wide screen replicate 1 and replicate 2.

**Table S2:** Results for each gene in the genome-wide screen, merged across sgRNAs and replicates. Outputs are included for our pipeline, MAGeCK, and casTLE.

**Table S3:** sgRNA sequences for custom CRISPR mini-pools, including the top hit mini-pool, SWI/SNF mini-pool, and perturb-seq micropool.

**Table S4:** sgRNA counts for all samples in the *in vitro* and *in vivo* mini-pool screens.

**Table S5:** z-score and FDR for each gene in the *in vitro* and *in vivo* mini-pool screens, merged across sgRNAs and replicates.

**Table S6:** sgRNA counts for all samples in the *in vivo* SWI/SNF mini-pool screens.

**Table S7:** z-score and FDR for each gene in the *in vivo* SWI/SNF mini-pool screens, merged across sgRNAs and replicates.

**Table S8:** sgRNA sequences and counts for all samples in the *in vivo* human mini-pool screen.

---

**Figure S1:**
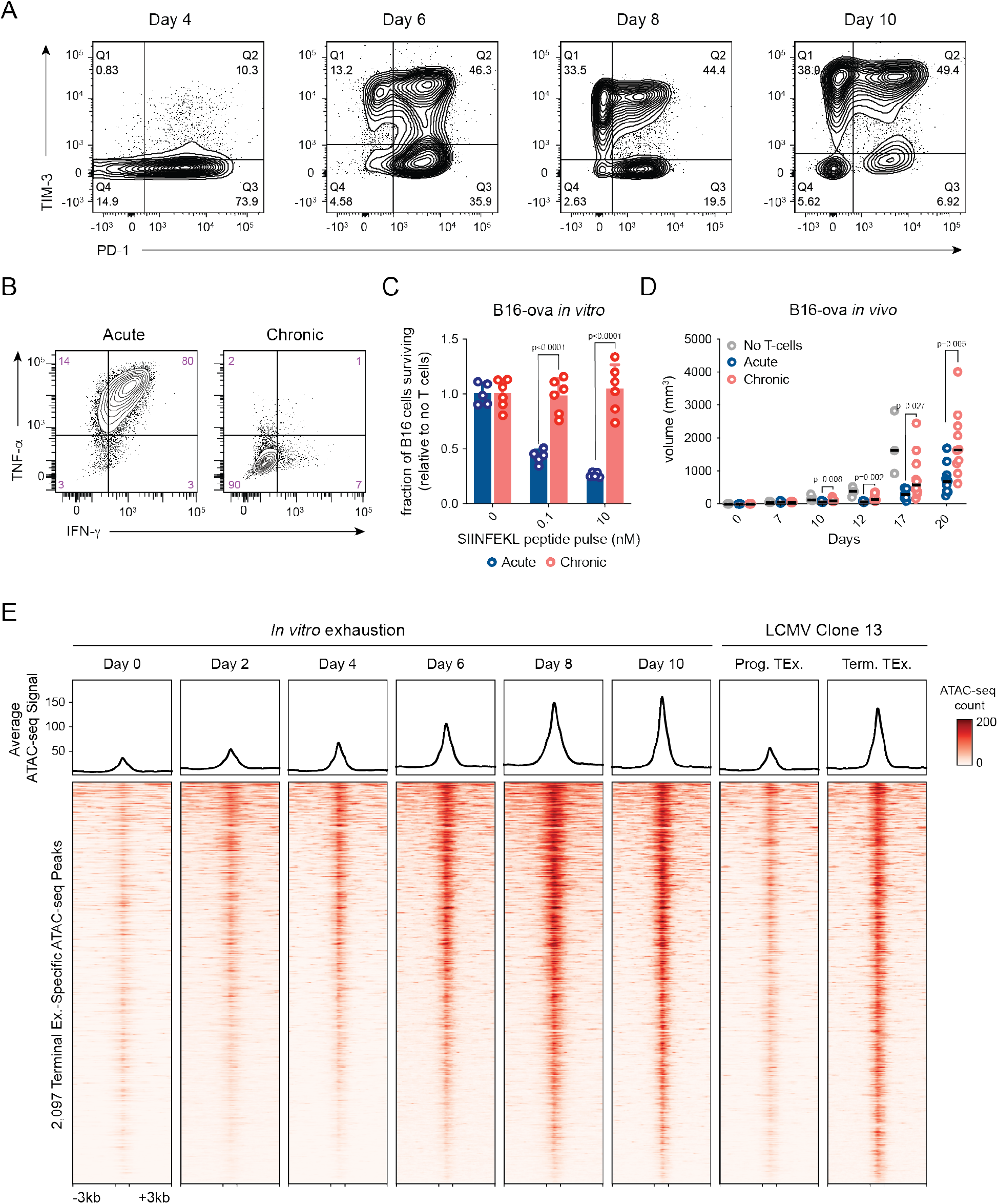
Additional characterization of in vitro assay. **(A)** Surface phenotype of chronically stimulated T cells throughout the *in vitro* exhaustion assay. **(B)** Effector cytokine production of acutely (left) and chronically (right) stimulated T cells. Cells were restimulated with PMA and ionomycin 8 days after initial stimulation. **(C)** Survival of B16 cells after co-culture with acutely or chronically stimulated OT-1 T cells. Tumor cells were pulsed with cognate peptide (SIINFEKL). **(D)** B16-ovalbumin tumor growth *in vivo* after adoptive transplant of acutely or chronically stimulated T cells. **(E)** Heatmap showing ATAC-seq coverage of each peak in the “Terminal Exhaustion peak set” for each time point in the in vitro exhaustion assay. Reference data from T cells in LCMV is also included.

**Figure S2:**
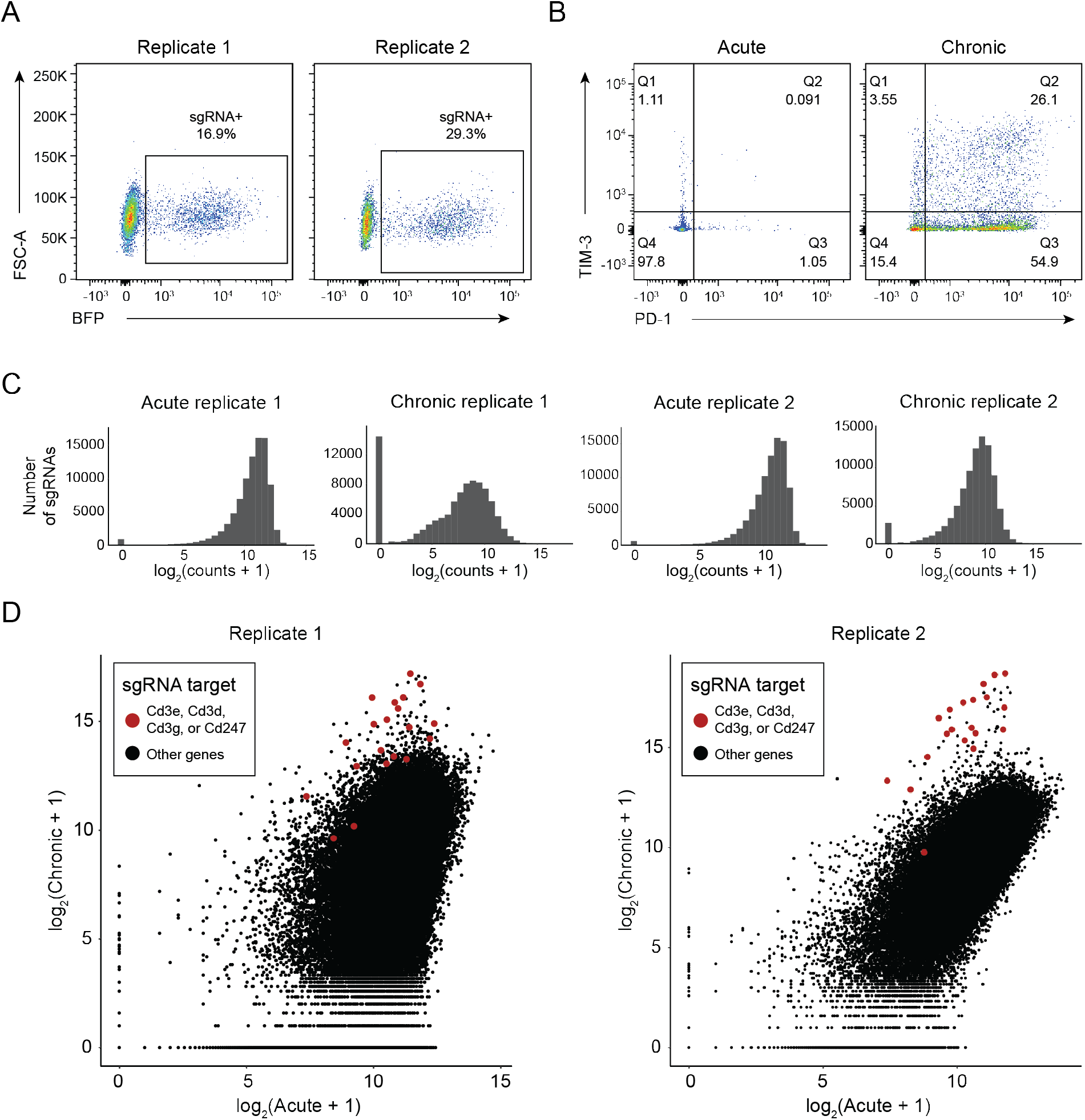
Quality control data for *in vitro* genome wide screen. **(A)** Expression of BFP on day 2 of the screen. **(B)** Surface phenotype of cells before gDNA extraction. **(C)** sgRNA representation of each sample. **(D)** sgRNA count correlations (Acute vs Chronic) for each replicate. CD3 subunits are shown in red, all other sgRNAs in black.

**Figure S3:**
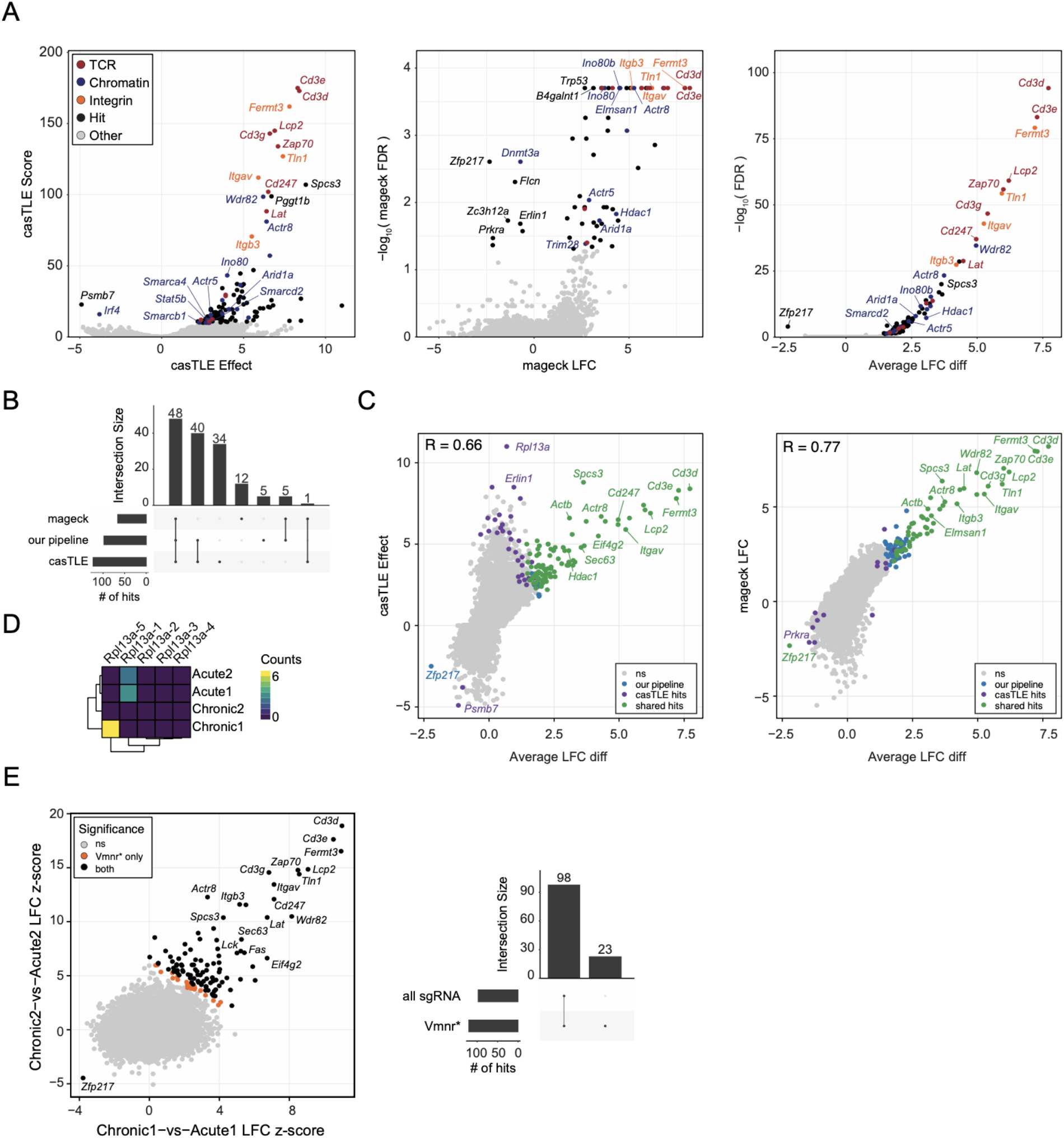
Comparison of CRISPR analysis strategies. **(A)** Volcano plots of genome wide CRISPR screen results using casTLE (left), MAGeCK (center), and our pipeline (right). **(B)** Comparison of hit lists for each of the three pipelines. **(C)** Comparison of LFC difference computed by our pipeline to the casTLE Effect (left) and MAGeCK LFC (right). **(D)** Counts table shown for *Rpl13a*. **(E)** Genome wide screen results when z-scores are computed relative to all sgRNAs or a set of olfactory receptors (Vmnr* genes).

**Figure S4:**
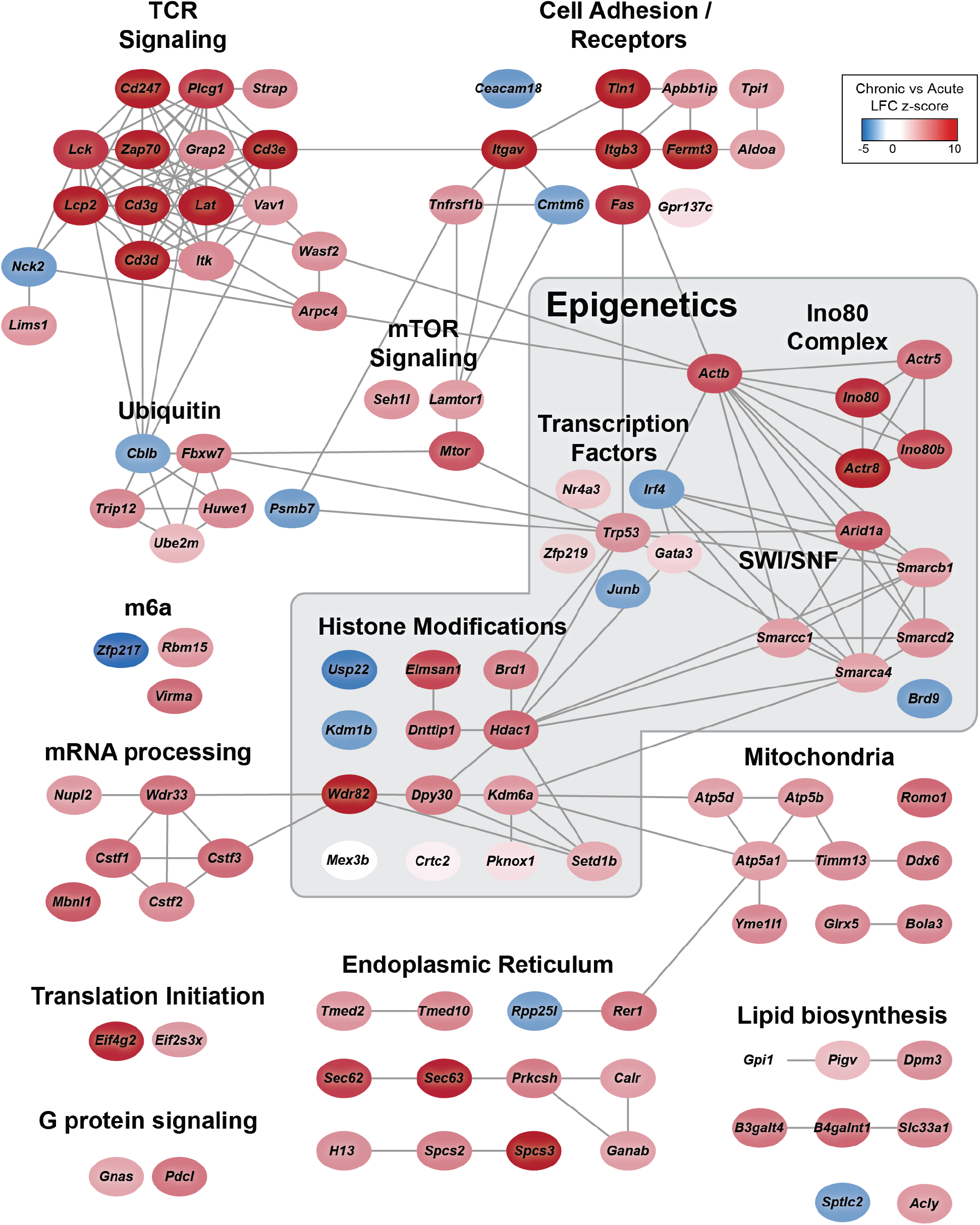
Cytoscape network representation of top hits. Top positive and negative hits from the genome-wide screen are shown. Each protein is represented by a node in the cytoscape network, colored by its z-score in the genome-wide screen. Nodes are connected if there is a high confidence protein-protein interaction in the string-db database (Szklarczyk et al., 2019).

**Figure S5:**
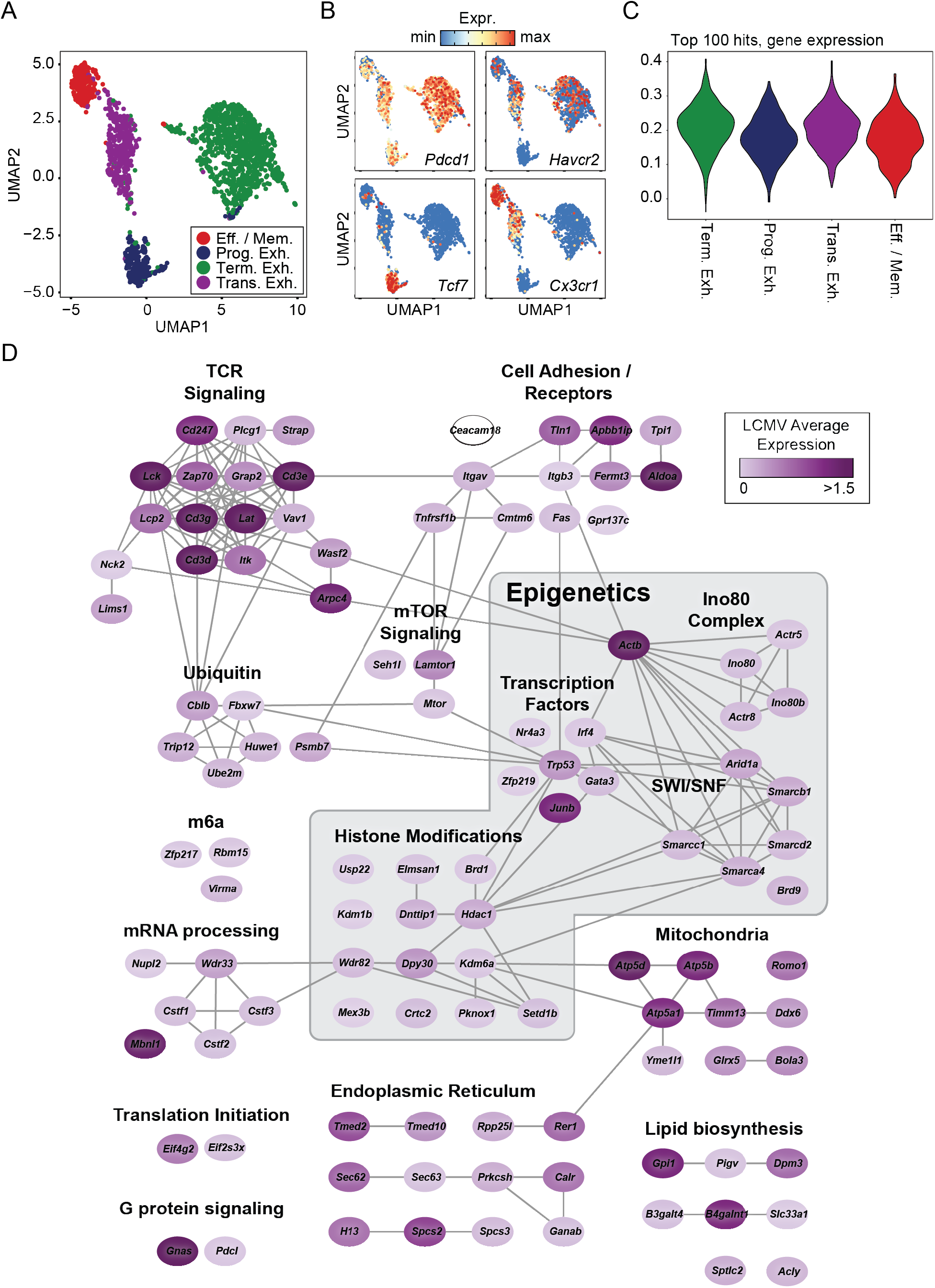
LCMV Clone 13 expression analysis of top hits. **(A)** Cell types identified in previously published scRNA-seq data (Raju et al., 2021). **(B)** Expression of *Pdcd1*, *Havcr2*, *Tcf7*, and *Cx3cr1* in single cells. **(C)** Expression of the gene module containing the top 100 *in vitro* hits across clusters. **(D)** Cytoscape network of top hits colored by average expression across all single cells.

**Figure S6:**
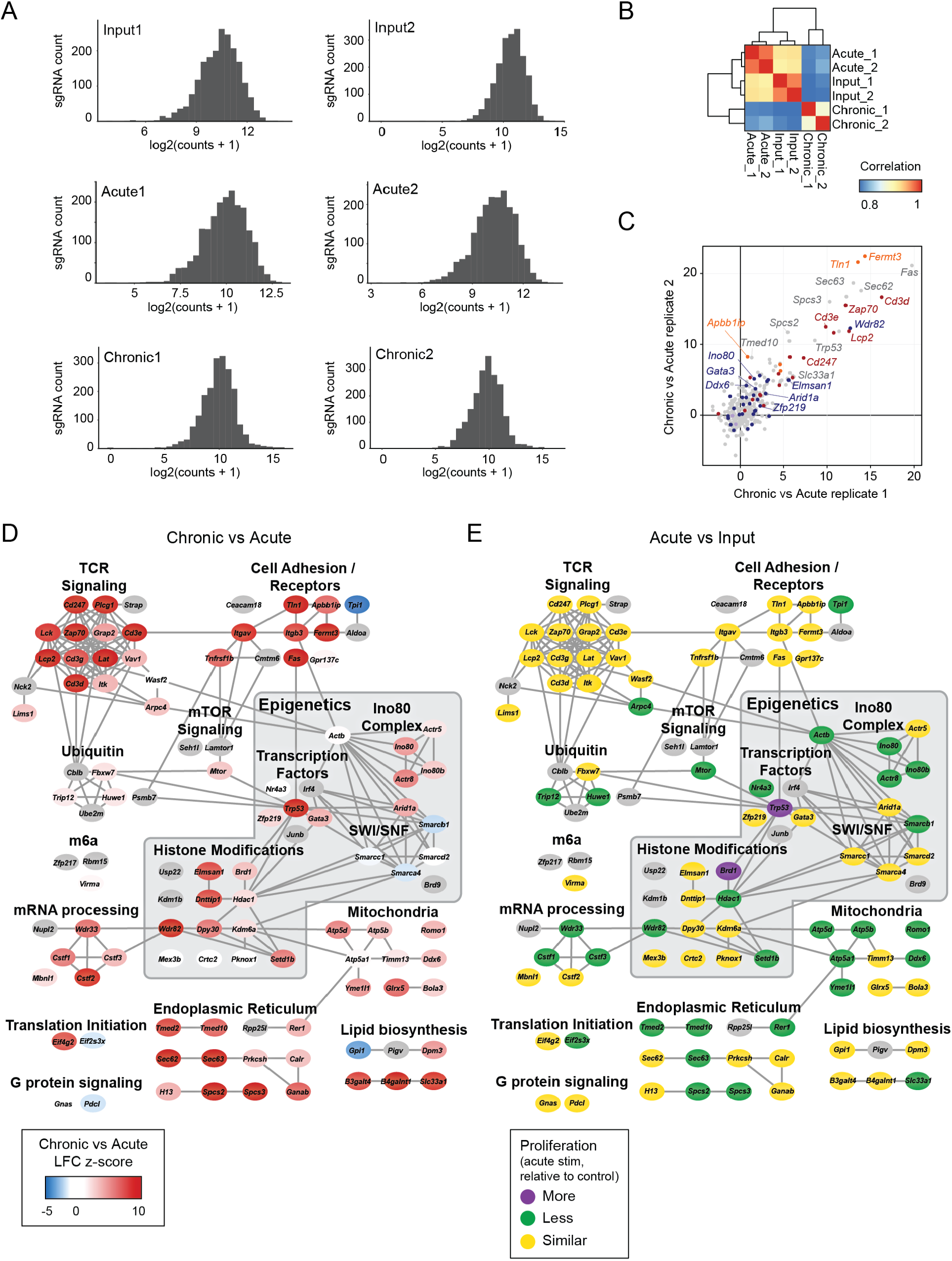
Additional data for targeted *in vitro* screening. **(A)** sgRNA representation of each sample in the *in vitro* mini-pool screen. **(B)** Correlation of the sgRNA counts of each sample in the mini-pool screen. **(C)** Correlation of the Chronic vs Acute replicate z-scores. **(D)** Cytoscape interaction network with genes colored by their z-score in the Chronic vs Acute mini-pool screen. **(E)** Cytoscape interaction network with genes colored by their fitness categorization in acute stimulation.

**Figure S7:**
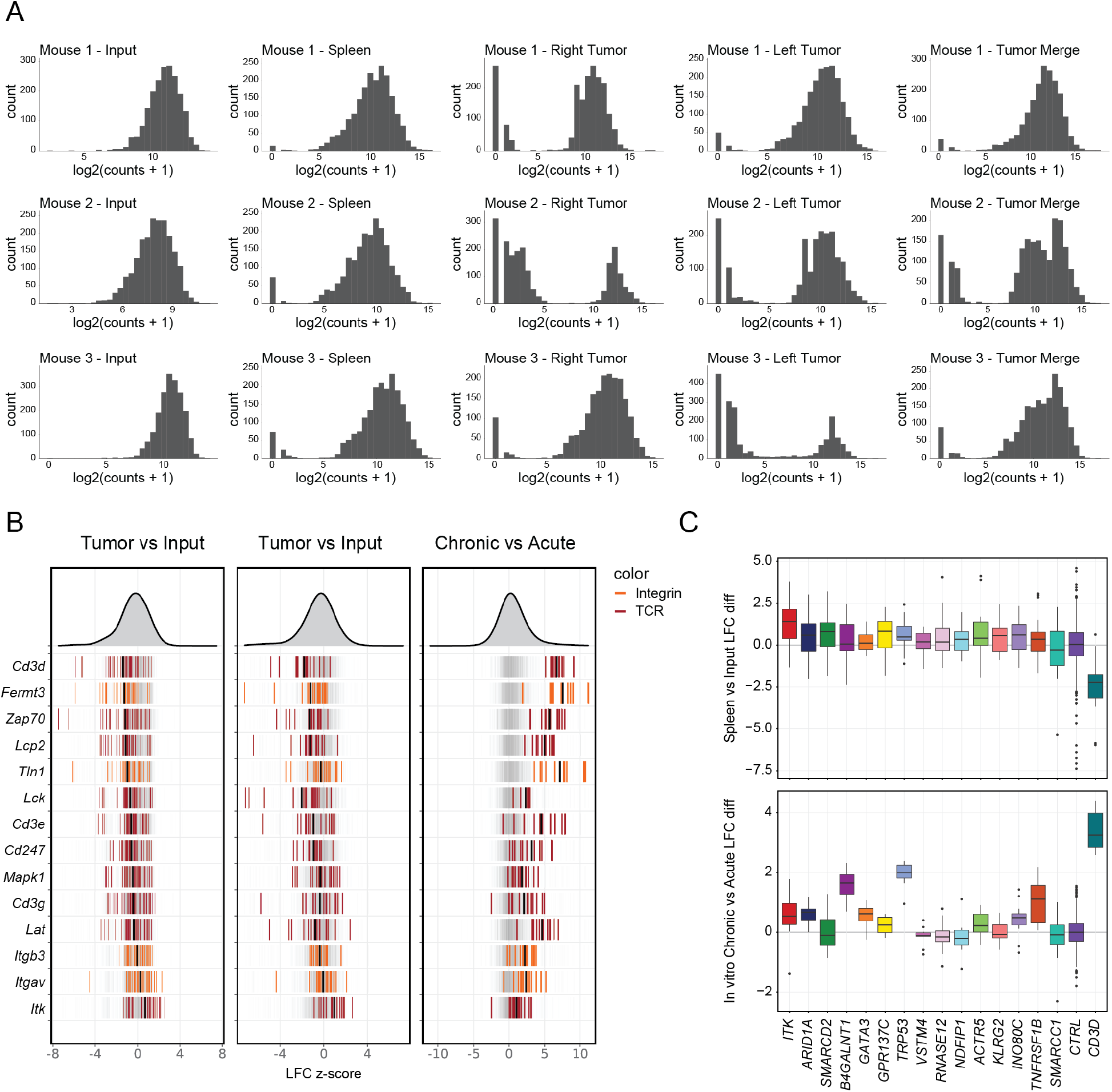
Additional data for targeted *in vivo* screening. **(A)** sgRNA pool coverage for each sample in the *in vivo* mini-pool screen. **(B)** sgRNA residuals in tumors, spleens, and *in vitro* mini-pool Chronic vs Acute for selected genes in the “TCR signaling” and “Integrin signaling” categories. **(C)** Boxplot of spleen vs input and acute vs chronic log fold change for each sgRNA targeting the indicated gene, with the mean control log fold change subtracted.

**Figure S8:**
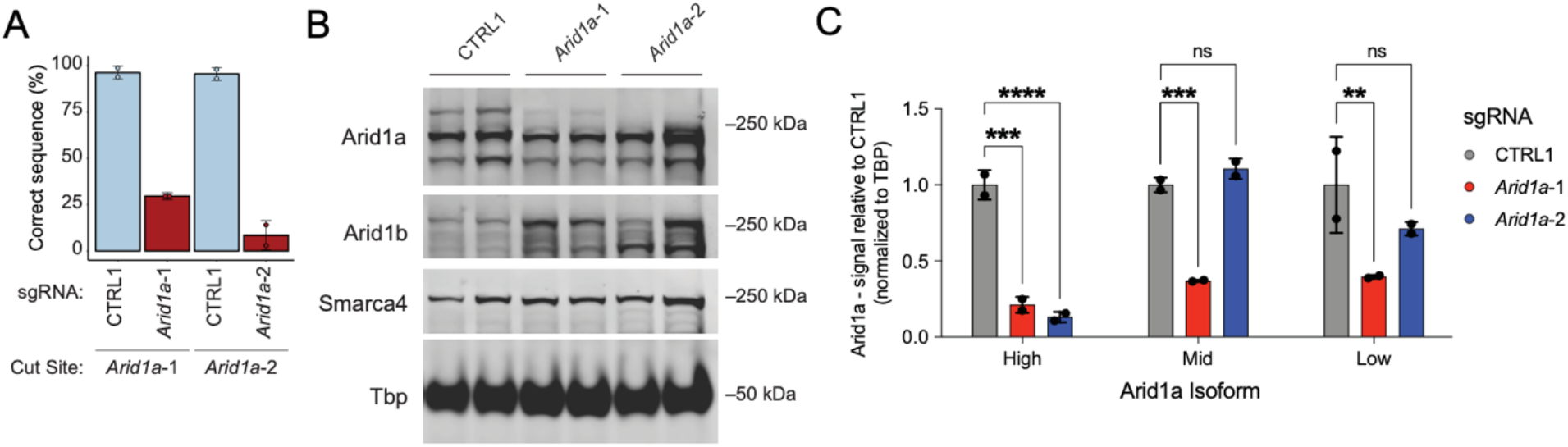
Validation of *Arid1a*-targeting sgRNAs. **(A)** Sanger sequencing (TIDE) analysis of editing efficiency of *Arid1a* sgRNAs. **(B)** Western blot analysis of protein knockdown for Arid1a sgRNAs, as well as Arid1b and Smarca4 expression. **(C)** Quantification of protein knockdown for each identified isoform of *Arid1a* (panel C three bands). * p < 0.05, ** p < 0.01, *** p < 0.001, **** p < 0.0001.

**Figure S9:**
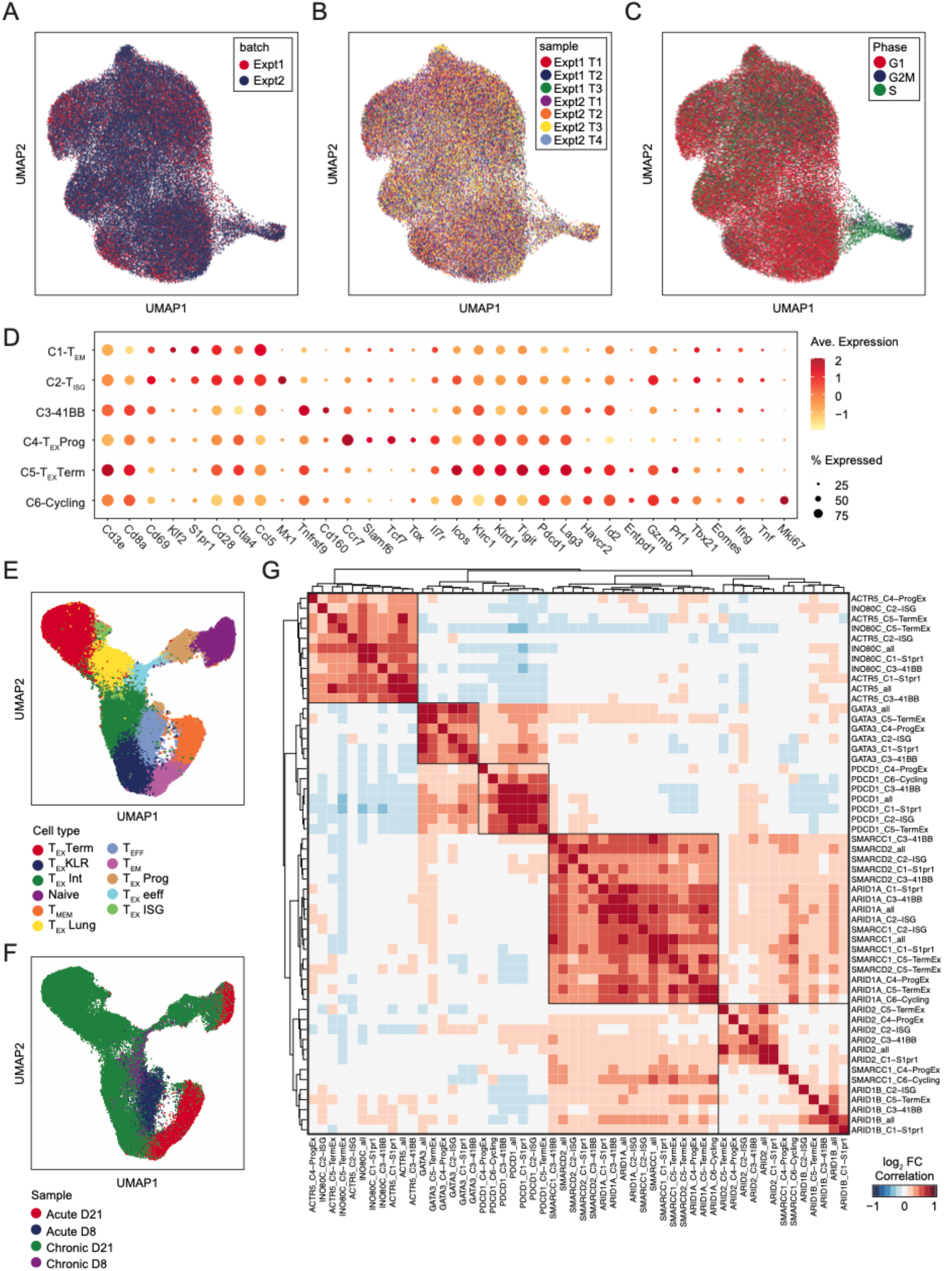
Additional data on the *in vivo* Perturb-seq experiment. **(A)** scRNA-seq profiles of TILs colored by each independent experiment. **(B)** scRNA-seq profiles of TILs colored by each sample. **(C)** scRNA-seq profiles of TILs colored by predicted phase of the cell cycle. **(D)** Additional marker genes shown for each cluster. **(E)** Expanded reference LCMV dataset with single cell profiles colored by LCMV cluster. Data from (Daniel et al., 2021). **(F)** Expanded LCMV dataset with single cell profiles colored by LCMV infection (Acute corresponds to Armstrong infection while Chronic corresponds to Clone 13) and time point (Day 8 or Day 21 post infection). **(G)** Heatmap of the correlation of gene expression differences subsetted on each cluster. The indicated gene knockdown was compared to CTRL1 cells within each cluster. Comparisons with <150 cells in the comparison groups are excluded due to lack of statistical power.

**Figure S10:**
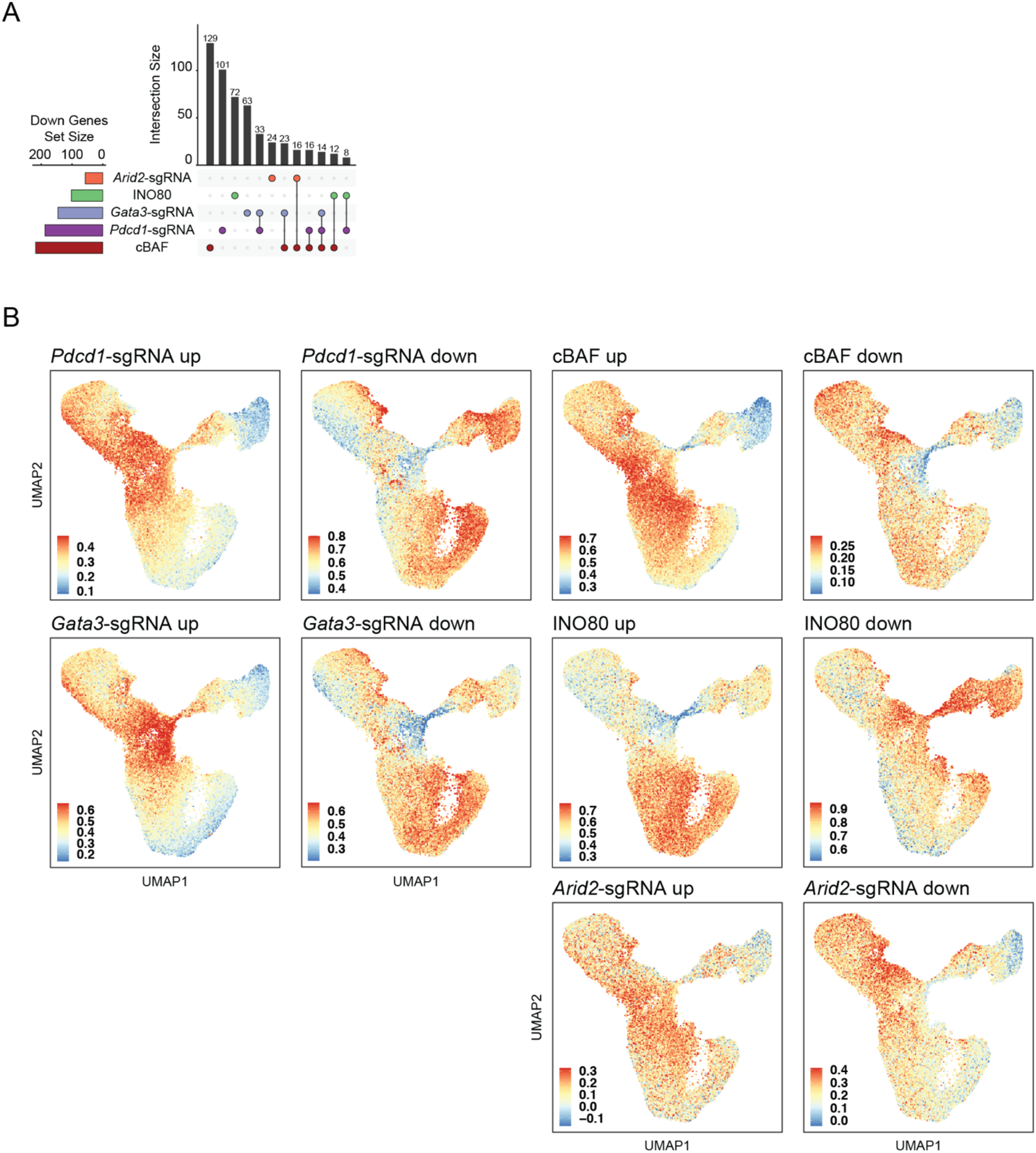
Additional data on up- and downregulated gene sets. **(A)** Comparison of gene sets downregulated by perturbation of cBAF subunits, INO80 subunits, or *Pdcd1*-sgRNA, *Gata3*-sgRNA, or *Arid2*-sgRNA. **(B)** Module scores of the indicated gene sets computed for each cell in the expanded LCMV reference dataset.

## Materials and Methods

### Mice

All mice were procured from JAX. Wild type mice were C57BL/6J mice (JAX: 000664). Rosa26-Cas9 knockin mice were bred in house (JAX: 026179). OT-1 mice (JAX: 003831) were crossed with Cas9 mice and then bred in house. Rag1^-/-^ mice were bred in house (JAX: 002216). C57BL/6 scid mice (JAX: 001913) and NSG mice (JAX: 005557) were procured from JAX.

### Primary murine T cell isolation and culture

Spleens were collected and mashed through a 70 uM filter. Red blood cells were lysed with ACK lysis buffer (Gibco) and incubated for 6 mins before washing with PBS. Cells were counted and then resuspended in MACS buffer (PBS + 0.5% BSA + 2 μM EDTA). CD8 T cells were enriched using the mouse CD8^+^ T cells isolation kit from Miltenyi (Miltenyi Cat# 130-104-075) and then resuspended in RPMI with 10% FBS, 1 % Sodium pyruvate, 1% Non-essential amino-acids, 100U Pen/Strep, 50 nM of B-mercaptoethanol (cRPMI) and supplemented with 10 ng/ml of mouse IL-2. Cells were seeded at a concentration of 1 million cells/ml on plates coated with 5 ug/ml of anti-CD3 and 2 ug/ml of anti-CD28. Cells were kept on activation plates for 48 hours at the beginning of all experiments. CD8^+^ T cell purity was verified via flow cytometry. Cells were passaged every two days and maintained at 1 million cells/mL.

### In vitro T cell exhaustion assay

To induce T cell exhaustion, chronic stimulation was performed using plates coated with anti-CD3 at 5 ug / mL. Cells were passaged onto a fresh coated plate every two days. Activation and T cell media are described above.

### Measurement of cytokine production

T cells were re-stimulated with phorbol myristate acetate (Sigma, 50 ng/mL) and ionomycin (Sigma, 500 ng/mL). After 90 minutes, cells were treated with brefeldin A to block cytokine secretion. Then, 3 hours later, cells were stained for surface markers and simultaneously labeled with Live/Dead Blue Viability Dye (Thermo Fisher) for 20 min at 4 °C. Cells were washed twice and fixed overnight using a FoxP3 Fixation/Permeabilization Kit (Thermo Fisher). The following day, cells were washed and stained for intracellular cytokines at room temperature for 1 hour. They were then washed three times and analyzed using an LSR Fortessa machine (Beckman Dickinson). Analysis of mean fluorescence intensity was performed using FlowJo v.10.0. All experiments were performed at least two independent times. Antibodies used (at 1:100 unless otherwise noted) were TNF-PE (BioLegend, MP6-XT22, 506306), PD-1-PECy7 (BioLegend, RMP1–30, 109110) IFN-γ-FITC (BioLegend, XMG1.2, 505806), CD4-BV711 (BioLegend, RM4– 5, 100550), and CD8α-BV786 (BioLegend, 53–6.7, 100750).

### Growth curves

After activation (described above), T cells were plated in 24-well plates at 5 × 10^5^ cells in 1 mL of RPMI-1640 medium containing 10% FBS, 2 mM l-glutamine, 5 μM β-ME and 10 ng/mL IL-2, and with (chronic) or without (acute) plate-bound anti-CD3. Every 2 days for the duration of the experiment, cells were collected, and cell number was counted using a Beckmann Coulter Counter with a cell volume gate of 75–4,000 femtoliters. Then, 50% of the cells were re-plated in 1 mL of fresh T cell medium. All experiments were performed at least two independent times.

### In vitro killing assay

B16 cells expressing a Luciferase reporter were pulsed with SIINFEKL peptide (Invivogen) at the concentrations noted in Figure S1 for 4 hours at 37C. They were then washed twice and plated at 4 x 10^4^ cells per well along with 1 × 10^5^ OT-1 transgenic T cells that had been acutely or chronically stimulated for 8 days as previously described. After 24 hours of co-culture, cells were lysed and luciferase activity was measured using a Luciferase Assay Kit (Promega) as per manufacturer’s instructions. Luciferase activity was normalized to cells cultured in the absence of T cells.

### B16-ovalbumin in vivo tumor models

C57BL/6 scid (Jackson 001913) mice were injected subcutaneously with 2 × 10^5^ B16-OVA cells in a 1:1 mix of PBS and Matrigel (Corning). 5 days later, 2 × 10^6^ OT-1 T cells that had been acutely or chronically stimulated as described previously were adoptively transferred to mice via retro-orbital injection. Mice were monitored daily and were killed for signs of morbidity.

### ATAC-seq sample processing and analysis

ATAC-seq was performed using the Omni-ATAC protocol (Corces et al., 2017). Briefly, 50,000 live cells were purified by flow cytometry immediately prior to ATAC-seq. Lysis, nuclei isolation, and transposition were performed according to the Omni-ATAC protocol. Libraries were prepared for sequencing and sequenced in 2×75 dual-indexed format on an Illumina NovaSeq.

Fastq files were trimmed using fastp (Chen et al., 2018) and aligned to the mm10 genome using hisat2 (Kim et al., 2019, p. 2). Reads were deduplicated and a bed file for each sample containing filtered, deduplicated ATAC-seq fragments was created. Peaks for each sample were called individually using MACS2 (Zhang et al., 2008) and then filtered into reproducible peaks based on peaks present in the majority of replicates for that sample. A union peak set for all samples was constructed by merging reproducible peaks for each sample into a set of high-confidence non-overlapping fixed width (500bp) peaks, which was used to create a peak by sample matrix used in downstream analysis. Differential peaks were determined using DESeq2 (Love et al., 2014, p. 2). Principal component analysis was performed on the peak matrix by first normalizing using ‘DESeq2::varianceStabilizingTransformation’ and then ‘stats::prcomp’. Genome track files were created by loading the fragments for each sample into R, and exporting bigwig files normalized by reads in transcription start sites using ‘rtracklayer::export’. Coverage files were visualized using the Integrative Genomics Viewer. For analysis of previously published ATAC-seq data (Miller et al., 2019), fastq files were downloaded from accession **GSE123236** and re-processed using our pipeline for consistency. For quantification of overlapping peaks between published data and in vitro assay data, a union peak set was created encompassing all samples and re-analyzed.

### Genome-wide sgRNA library

Retroviral Mouse Genome-wide CRISPR Knockout Library was a gift from Sarah Teichmann (Addgene #104861). The library was amplified via electroporation and confirmed by sequencing.

### sgRNA pool design and cloning

The sgRNA mini-pool was designed using our previously developed protocol for cloning into a lentiviral backbone and then subcloned into retroviral construct pMSCV (Flynn et al., 2021). The lentiCRISPR-v2 was a gift from Feng Zhang (Addgene plasmid #52961). pMSCV-U6sgRNA(BbsI)-PGKpuro2ABFP was a gift from Sarah Teichmann (Addgene plasmid #102796).

Briefly, six 20bp variable sgRNA sequences per target gene were obtained from the Broad Genetic Perturbation Platform (GPP) genome wide designs: sgRNA_design_10090_GRCm38_SpyoCas9_CRISPRko_NCBI_20200317.txt.gz, available online at https://portals.broadinstitute.org/gpp/public/dir?dirpath=sgrna_design. 100 non-targeting and 100 single-targeting negative control guides designed for the mouse genome, also from the Broad GPP web portal, were included. A “G” was added to the start of each 20bp sequence. This 21bp sequence was flanked by BsmBI-v2 enzyme sites and then two nested PCR handles. Pooled oligos were synthesized by Twist Bioscience. Oligos were amplified by two rounds of PCR and the lentiCRISPR-v2 backbone was digested overnight with Esp3I. One step digestion/ligation of amplified oligos into lentiCRISPR-v2 was performed at 37C for 1 hour in a 20 uL reaction with 1 uL T4 ligase, 1 uL Esp3I, 2 uL T4 ligase buffer, 200 ng digested backbone, and 50 ng amplified insert. Reaction was heat inactivated for 15 minutes at 65C and then 1 uL was electroporated using 25 uL Lucigen Endura electrocompetent cells and a BioRad MicroPulser with 0.1 cm gap cuvettes. After 1 hour recovery in SOC, a 1000x dilution was plated onto an agar plate to confirm library coverage. The remainder was cultured overnight in a 150 mL liquid culture and then purified by maxiprep. Finally, the pool was subcloned into pMSCV by Gibson Assembly of the sgRNA variable region amplified via PCR and pMSCV backbone pre-digested with BbsI. Electroporation was repeated as described above. Guide representation was confirmed by sequencing.

The sgRNA SWI/SNF mini-pool and micro-pool for Perturb-seq were designed with 4 guides per gene, as described above for the mini-pool using the Broad GPP mouse genome-wide designs. The SWI/SNF mini-pool contained 50 single-targeting controls and Perturb-seq micro-pool contained 12 single-targeting controls. Two primers were ordered per designed guide, for cloning via annealing. The pMSCV vector was digested with BbsI. All primer pairs were annealed separately. Annealed products were pooled equally, diluted, and then ligated into pMSCV. Amplification was performed using Stbl3 Chemically Competent cells (ThermoFisher C737303) and library coverage was confirmed via colony counting and then sequencing.

### Retrovirus production and transduction

The pMSCV plasmid was transfected into GP2-293 cells (Takara, RetroPack™ PT67 Cell Line) or 293T HEK cells at roughly 80% confluency in 15 cm tissue culture plates coated with poly-d-lysine. Viral supernatant was collected at 48h and 72h post-transfection, filtered via a 0.45 μm filtration unit (Millipore). Filtered virus was concentrated using the LentiX concentrator (Takara) at 1500 x g for 45 minutes. The concentrated supernatant was subsequently aliquoted, flash frozen, and stored in −80°C until use.

CD8^+^ T cells were transduced with concentrated retrovirus 24 hours after isolation. 4 ug/mL of polybrene was added to each well. Plates were sealed and then spun at 1100x g at 32 C for 90 minutes. 24 hours after spinfection (ie, starting on day 2) cells were checked for fluorescence via flow cytometry and 2 ug/mL puromycin was added to the media.

### sgRNA library preparation and sequencing

For samples from in vitro chronic culture, live cells were first isolated via FACS. gDNA was extracted using a commercially available kit (Zymo Cat# D3025). sgRNA libraries were prepared for sequencing as previously described (Flynn et al., 2021). Briefly, a standard three-step amplification protocol was used. First, sgRNAs were amplified off of gDNA using primers specific to the pMSCV vector for 22 cycles of PCR. 100 uL reactions with up to 4 ug of gDNA per reaction were used, and the number of reactions was scaled up until all gDNA was used. For sequencing of plasmid pools, this first PCR was skipped. For the second PCR, a 0-7bp offset was added to the front of the library using 8 pooled stagger primers to increase the diversity of the library. PCR2 primer target sites were nested inside those of PCR1 to improve the specificity of the product. Finally, in PCR3, index sequences were added. Libraries were sequenced in dual-indexed 1×75 bp or 1×150 bp format on either an Illumina NextSeq or NovaSeq.

### Bulk sgRNA screening data analysis

sgRNA sequencing data was analyzed using our previously published pipelines (Flynn et al., 2021). Briefly, fastq files were trimmed using ‘fastp -f 10 --max_len1=50’. Trimmed reads were aligned to a custom fasta file of the relevant pool (either the genome wide pool or the mini-pool) which was constructed by taking the sgRNA variable sequences and flanking them with the adjacent sequences in the pMSCV vector backbone. Alignment was performed using hisat2 with the --no-spliced-alignment option. Bam files were imported into R and converted into counts per guide using ‘Rsamtools::scanBam’. A table of guides per sample was constructed in R and normalized by multiplying each count by 1e6, dividing by the total counts in that sample, adding 1, and then log2 normalizing. Log fold changes between two conditions (chronic vs acute or tumor vs input) were computed and then z-scored by subtracting the reference LFC average and dividing by the reference LFC standard deviation. For genome-wide screens, all guides were used as the reference and for mini-pool screens the control guides were used as the reference. P-values were computed from z-scores using the normal distribution and then FDR was computed by correcting for multiple hypothesis testing using ‘p.adjust’ in R.

### GO Term analysis

For gene categorizations shown in Figure 2B and elsewhere, gene sets were defined as: TCR - KEGG_T_CELL_RECEPTOR_SIGNALING_PATHWAY, Chromatin - GOCC_CHROMATIN, Integrin - GOBP_INTEGRIN_ACTIVATION, Inhibitory receptor - GOBP_NEGATIVE_REGULATION_OF_LYMPHOCYTE_ACTIVATION. Gene lists were manually supplemented with the following genes: Chromatin - *ZFP219, TBX21, KDM6A, ELMSAN1, DNTTIP1, SETD1B, TADA2B, ZFP217, EOMES*. Integrins - *ITGB3, APBB1IP, ITGAV*. Inhibitory receptors - *PDCD1.* For the gene set enrichment analysis shown in Figure 2D and elsewhere, the indicated gene list was uploaded to the online gProfiler tool (available at https://biit.cs.ut.ee/gprofiler/gost).

### Cytoscape interaction network

100 top enriched genes and 20 top depleted genes were imported into Cytoscape (Shannon et al., 2003). Edges were created by using the stringApp Cytoscape plugin to import known protein-protein interactions curated from string-db (Szklarczyk et al., 2019). A cutoff of stringdb score ≥ 0.75 was used to filter these protein protein interactions, which represents a conservative cutoff for identifying only high confidence interactions. Nodes were grouped based on GO Term analysis, subcellular localization, and/or manual curation. A small number of poorly characterized and/or disconnected nodes were removed from the visualization.

### Tumor inoculation and T cell adoptive transfer for in vivo CRISPR experiments

MC-38 or B16 cells ectopically expressing an mCherry-ovalbumin fusion construct were prepared for injection by resuspending in a 1:1 mixture of matrigel and PBS. 10^6^ cells per tumor were injected subcutaneously into the flanks of Rag1^-/-^ mice (two tumors per mouse). Tumors were measured every three days. Cas9/OT-1 CD8^+^ T cells were transduced with sgRNA pools or individual sgRNAs and selected with puromycin for 4 days, as described above. T cells were then intravenously injected into tumor-bearing mice on day 6. For *in vivo* competition assays, cells were mixed immediately prior to injection. 9 days after T cell injection, the spleen and tumors were harvested from each mouse.

### Tissue Processing and Isolation of Tumor Infiltrating Lymphocytes

Tumors were weighed and then minced into small pieces. The tumors were transferred to a gentleMACS C tube and digested in the protocol-recommended enzyme mix with a gentleMACS octo dissociator using the soft/medium tumor program. Tumor suspensions were then filtered with a 70 uM filter and then subject to RBC lysis. Spleens were mashed and filtered through a 70 uM strainer, then treated with RBC lysis buffer. For bulk sgRNA sequencing and Perturb-seq, T cells were isolated from the tumors and/or spleens by FACS. Samples were washed twice with MACS buffer and stained for 30 minutes on ice. CD8^+^ BFP^+^ cells were isolated via flow cytometry.

### Competition assay for validation of individual sgRNA proliferation

The pMSCV retroviral vector was modified to replace the BFP-puromycin fusion with a VEX-puromycin fusion. Individual guides were cloned by annealing pairs of primers, as described above. The *Arid1a*-1 sgRNA sequence used was GCAGCTGCGAAGATATCGGG and the *Arid1a*-2 sequence used was CAGCAGAACTCGCACGACCA. The CTRL sgRNA sequence used was CTTACTCGACGAATGAGCCC. Tumor processing was performed as described above for the *in vivo* validation.

### Validation of Arid1a-targeting sgRNAs

Tracking of indels by decomposition (TIDE): Genomic DNA was isolated from transduced cells using a commercially available kit (Zymo Cat# D3025). PCR reactions were performed with primers surrounding the expected edit site and 50 ng of input DNA. PCR conditions were 30 seconds at 98C, followed by 10 seconds at 98 C, 10 seconds annealing at 60C, 25 seconds at 72C for 35 cycles, then 2 minutes at 72C. The PCR amplicons were purified with a commercially available Zymo DNA clean up kit and sanger sequenced. Quantification of edits was performed using the online tool https://tide.nki.nl/.

Western blot: Protein lysates were prepared from mouse T cells transduced with the indicated sgRNA using a radioimmunoprecipitation assay (RIPA) buffer system (Santa Cruz, sc-24948). Protein concentrations were quantified using the bicinchoninic Acid (BCA) assay (Pierce, ThermoFisher 23225). 20 µg of protein per sample was loaded and run on a 4–12% Bis-Tris PAGE gel (NuPAGE 4-12% Bis-Tris Protein Gel, Invitrogen) and transferred onto a polyvinylidene fluoride (PVDF) membrane (Immobilon-FL, EMD Millipore). Membranes were blocked with 5% milk in PBST for 1 h at room temperature (RT) and incubated with primary antibodies against Arid1a (rabbit, 1:1000, Cell Signaling, 12354S: Lot 4), Arid1b (mouse, 1:1000, Abcam, ab57461: Lot GR3345290-4), Smarca4 (rabbit, 1:1000, Cell Signaling, 49360S: Lot 3) and Tbp (mouse, Abcam, ab51841: Lot GR3313213-3) overnight at 4 °C. Membranes were washed three times with PBST and then incubated with near-infrared fluorophore-conjugated species-specific secondary antibodies: Goat Anti-Mouse IgG Polyclonal Antibody (IRDye 680RD, 1:10,000, LI-COR Biosciences, 926-68070) or Goat Anti-Rabbit IgG Polyclonal Antibody (IRDye 800CW, 1:10,000, LI-COR Biosciences, 926-32211) for 1 hour at RT. Following secondary antibody application, membranes were washed three times with PBST, and then imaged using a LI-COR Odyssey CLx imaging system (LI-COR). Protein band intensities were quantified using Image Studio Lite (LI-COR) with built-in background correction and normalization to Tbp controls. Statistical analysis comparing Arid1a levels normalized to Tbp was performed using Dunnett’s multiple comparisons test on Prism (v9.2.0).

### In vitro experiments in primary human T cells

T cell expansion and viability assays: T cells were activated for 4 days at a 1:3 ratio of T cells to anti-CD3/28 Dynabeads (Invitrogen). T cell expansion assays were performed with IL-2 in the culture medium at 10 ng/mL. Cell counts and viability measurements were obtained using the Cellaca Mx Automated Cell Counter (Nexcelom). Cells were stained with acridine orange and propidium iodide to assess viability.

Targeted CRISPR gene editing: Ribonucleoprotein (RNP) was preparing using synthetic sgRNA with 2’-O-methyl phosphorothioate modification (Synthego) diluted in TE buffer at 100 µM. 5 µl sgRNA was incubated with 2.5 µl Duplex Buffer (IDT) and 2.5 µg Alt-R S.p. Cas9 Nuclease V3 (IDT) for 30 minutes at room temperature. 100 µl reactions were assembled with 10 million T cells, 90 µl P3 buffer (Lonza), and 10 µl RNP. Cell were pulsed with protocol EO115 using the P3 Primary Cell 4D-Nucleofector Kit and 4D Nucleofector System (Lonza). Cells were recovered immediately with warm media for 6 hours. Guide sequences: AAVS1-sg1 5’ GGGGCCACUAGGGACAGGAU 3’, ARID1A-sg58 5’ CCUGUUGACCAUACCCGCUG 3’, ARID1A-sg60 5’ UGUGGCUGCUGCUGAUACGA 3’.

Assessment of targeted CRISPR gene editing: 4-7 days after editing, genomic DNA was extracted with QuickExtract DNA Extraction Solution (Lucigen) and ∼500 bp regions flanking the cut site were amplified with Phusion Hot Start Flex 2X Master Mix (New England Biolabs) according to manufacturer’s instructions. Sanger sequencing traces were analyzed by Inference of CRISPR Editing (ICE).

### Pooled CRISPR screen in primary human T cells in vivo

Activated human T cells from two donors were transduced by lentivirus to express the NY-ESO specific TCR, in parallel to lentiviral transduction of a sgRNA library. 24 hours after transduction, cells were electroporated with Cas9 Protein, as previously described (Shifrut et al., 2018). After electroporation, T cells were expanded in complete X-vivo 15 medium and split every two days, supplementing IL-2 at 50 U/ml. On Day 7, 2 NSG mice per donor were injected subcutaneously with 1 x 10^6^ A375 cells, as previously described (Roth et al, Cell 2020). 1 x 10^6^ TCR-positive T cells were transferred to mice 7 days later via retro-orbital injection. Tumors and spleens were collected 7 days after T cell transfer and processed to single cell suspension, as described previously (Roth et al., 2020). T cells were sorted by CD45 staining and gDNA was extracted using commercial kits. Library preparation, next generation sequencing and analysis was performed as previously described (Shifrut et al., 2018). The guide abundance in the spleen and tumor of each mouse was used to calculate log fold change of each guide, and MAGeCK scores were calculated with default parameters.

### Direct-capture Perturb-seq

For Perturb-seq experiments, we used direct-capture Perturb-seq because it does not require a vector with a barcode sequence separate from the sgRNA, or other modifications to standard sgRNA vectors, and thus was immediately compatible with our retroviral reagents (Replogle et al., 2020). We adapted the 10x Chromium Next GEM Single Cell V(D)J Reagent Kits v1.1 5’ scRNA with Feature Barcoding reagents and protocol to be compatible with direct capture of sgRNAs in single cells. Our procedure is conceptually similar to that of Replogle et al and the modifications to the 10X genomics protocol are summarized here. For Step 1, GEM Generation and Barcoding, 5 pmol of primer KP_bead_sgRNA_RT was spiked into the reaction, enabling capture of sgRNAs in droplets and then reverse transcription of sgRNAs. Step 3.2B, Supernatant Cleanup for Cell Surface Protein Library was performed to isolate the sgRNA library. Finally, 2 uL of the product of Step 3.2B was amplified and indexed using 3 rounds of PCR. The 250bp library was purified via agarose gel and sequenced together with the gene expression (GEX) library in 26×91 format, according to 10X protocol guidelines.

Fastq files were processed using the 10X cellranger count pipeline with feature barcode analysis enabled to process the GEX library and sgRNA library together. The mm10 reference transcriptome was used for the GEX library. For the sgRNA library, a feature reference spreadsheet was constructed which contained the variable sequence of each guide (reverse complemented since it was sequenced as part of read 2), guide ID, and target gene. The filtered matrices for both ‘Gene Expression’ and ‘CRISPR Guide Capturè were loaded into Seurat for downstream analysis (Hao et al., 2021). The Seurat ‘IntegrateDatà utility was used to merge the samples from the two independent experiments.

To assign sgRNAs to cells, we computed row z-scores for the ‘CRISPR Guide Capturè matrix. We computed z-scores quantifying how enriched each sgRNA was relative to other sgRNAs detected in the same cell. We also computed the difference in z-scores between the most-enriched and second-most enriched sgRNA. Cells which had a maximum sgRNA z-score ≥ 5 and a z-score difference ≥ 2 was determined to contain the guide with maximum z-score, while cells with no sgRNA counts were assigned as “no guide,” and other cells were assigned “multi guide”. The guide assignments were added to the Seurat metadata for downstream processing. Seurat cell cycle scoring was used to predict the cell cycle phase of each single cell. For volcano plot analysis, significantly differential genes were identified as FDR < 0.05. For comparisons of different gene sets across perturbations, an addition fold change cutoff was applied of average log_2_ FC > 0.1 or average log_2_ FC < −0.1. For categorization of shared ‘up’ and ‘down’ gene sets within the cBAF and INO80 complexes (analysis shown in Figure 7D-E), the union set of significantly differential genes within each complex was aggregated, and then ‘up’ and ‘down’ genes for each subunit were defined simply as LFC > 0 or LFC < 0. This strategy was chosen to compare gene sets despite the different amounts of cells collected for each perturbation and resulting difference in statistical power to reach the FDR < 0.05 threshold. Seurat gene module scoring was used to convert the LCMV gene sets (consisting of the top 100 marker genes per LCMV cluster) into a gene module score for each cell in the perturb-seq dataset. Gene module scoring was also used to convert the upregulated and downregulated gene sets into module scores for each cell in the expanded LCMV data set, as shown in Figure S10.

## References

Adamson, B., Norman, T.M., Jost, M., Cho, M.Y., Nuñez, J.K., Chen, Y., Villalta, J.E., Gilbert, L.A., Horlbeck, M.A., Hein, M.Y., Pak, R.A., Gray, A.N., Gross, C.A., Dixit, A., Parnas, O., Regev, A., Weissman, J.S., 2016. A Multiplexed Single-Cell CRISPR Screening Platform Enables Systematic Dissection of the Unfolded Protein Response. Cell 167, 1867–1882.e21. https://doi.org/10.1016/j.cell.2016.11.048

Alfei, F., Kanev, K., Hofmann, M., Wu, M., Ghoneim, H.E., Roelli, P., Utzschneider, D.T., von Hoesslin, M., Cullen, J.G., Fan, Y., Eisenberg, V., Wohlleber, D., Steiger, K., Merkler, D., Delorenzi, M., Knolle, P.A., Cohen, C.J., Thimme, R., Youngblood, B., Zehn, D., 2019. TOX reinforces the phenotype and longevity of exhausted T cells in chronic viral infection. Nature 571, 265–269. https://doi.org/10.1038/s41586-019-1326-9

Ataide, M.A., Komander, K., Knöpper, K., Peters, A.E., Wu, H., Eickhoff, S., Gogishvili, T., Weber, J., Grafen, A., Kallies, A., Garbi, N., Einsele, H., Hudecek, M., Gasteiger, G., Hölzel, M., Vaeth, M., Kastenmüller, W., 2020. BATF3 programs CD8+ T cell memory. Nat Immunol 21, 1397–1407. https://doi.org/10.1038/s41590-020-0786-2

Barber, D.L., Wherry, E.J., Masopust, D., Zhu, B., Allison, J.P., Sharpe, A.H., Freeman, G.J., Ahmed, R., 2006. Restoring function in exhausted CD8 T cells during chronic viral infection. Nature 439, 682–687. https://doi.org/10.1038/nature04444

Beltra, J.-C., Manne, S., Abdel-Hakeem, M.S., Kurachi, M., Giles, J.R., Chen, Z., Casella, V., Ngiow, S.F., Khan, O., Huang, Y.J., Yan, P., Nzingha, K., Xu, W., Amaravadi, R.K., Xu, X., Karakousis, G.C., Mitchell, T.C., Schuchter, L.M., Huang, A.C., Wherry, E.J., 2020. Developmental Relationships of Four Exhausted CD8+ T Cell Subsets Reveals Underlying Transcriptional and Epigenetic Landscape Control Mechanisms. Immunity 52, 825–841.e8. https://doi.org/10.1016/j.immuni.2020.04.014

Chen, J., López-Moyado, I.F., Seo, H., Lio, C.-W.J., Hempleman, L.J., Sekiya, T., Yoshimura, A., Scott-Browne, J.P., Rao, A., 2019. NR4A transcription factors limit CAR T cell function in solid tumours. Nature 567, 530–534. https://doi.org/10.1038/s41586-019-0985-x

Chen, S., Zhou, Y., Chen, Y., Gu, J., 2018. fastp: an ultra-fast all-in-one FASTQ preprocessor. Bioinformatics 34, i884–i890. https://doi.org/10.1093/bioinformatics/bty560

Chen, Z., Arai, E., Khan, O., Zhang, Z., Ngiow, S.F., He, Y., Huang, H., Manne, S., Cao, Z., Baxter, A.E., Cai, Z., Freilich, E., Ali, M.A., Giles, J.R., Wu, J.E., Greenplate, A.R., Hakeem, M.A., Chen, Q., Kurachi, M., Nzingha, K., Ekshyyan, V., Mathew, D., Wen, Z., Speck, N.A., Battle, A., Berger, S.L., Wherry, E.J., Shi, J., 2021. In vivo CD8+ T cell CRISPR screening reveals control by Fli1 in infection and cancer. Cell 184, 1262–1280.e22. https://doi.org/10.1016/j.cell.2021.02.019

Collier, J.L., Weiss, S.A., Pauken, K.E., Sen, D.R., Sharpe, A.H., 2021. Not-so-opposite ends of the spectrum: CD8+ T cell dysfunction across chronic infection, cancer and autoimmunity. Nat Immunol 22, 809–819. https://doi.org/10.1038/s41590-021-00949-7

Corces, M.R., Trevino, A.E., Hamilton, E.G., Greenside, P.G., Sinnott-Armstrong, N.A., Vesuna, S., Satpathy, A.T., Rubin, A.J., Montine, K.S., Wu, B., Kathiria, A., Cho, S.W., Mumbach, M.R., Carter, A.C., Kasowski, M., Orloff, L.A., Risca, V.I., Kundaje, A., Khavari, P.A., Montine, T.J., Greenleaf, W.J., Chang, H.Y., 2017. An improved ATAC-seq protocol reduces background and enables interrogation of frozen tissues. Nat Methods 14, 959– 962. https://doi.org/10.1038/nmeth.4396

Daniel, B., Yost, K.E., Sandor, K., Xia, Y., Qi, Y., Hiam-Galvez, K.J., Meier, S.L., Belk, J.A., Giles, J.R., Wherry, E.J., Chang, H.Y., Egawa, T., Satpathy, A.T., 2021. Divergent clonal differentiation trajectories of T cell exhaustion. https://doi.org/10.1101/2021.12.16.472900

DePeaux, K., Delgoffe, G.M., 2021. Metabolic barriers to cancer immunotherapy. Nat Rev Immunol. https://doi.org/10.1038/s41577-021-00541-y

Dixit, A., Parnas, O., Li, B., Chen, J., Fulco, C.P., Jerby-Arnon, L., Marjanovic, N.D., Dionne, D., Burks, T., Raychowdhury, R., Adamson, B., Norman, T.M., Lander, E.S., Weissman, J.S., Friedman, N., Regev, A., 2016. Perturb-Seq: Dissecting Molecular Circuits with Scalable Single-Cell RNA Profiling of Pooled Genetic Screens. Cell 167, 1853–1866.e17. https://doi.org/10.1016/j.cell.2016.11.038

Dong, M.B., Wang, G., Chow, R.D., Ye, L., Zhu, L., Dai, X., Park, J.J., Kim, H.R., Errami, Y., Guzman, C.D., Zhou, X., Chen, K.Y., Renauer, P.A., Du, Y., Shen, J., Lam, S.Z., Zhou, J.J., Lannin, D.R., Herbst, R.S., Chen, S., 2019. Systematic Immunotherapy Target Discovery Using Genome-Scale In Vivo CRISPR Screens in CD8 T Cells. Cell 178, 1189–1204.e23. https://doi.org/10.1016/j.cell.2019.07.044

Flynn, R.A., Belk, J.A., Qi, Y., Yasumoto, Y., Wei, J., Alfajaro, M.M., Shi, Q., Mumbach, M.R., Limaye, A., DeWeirdt, P.C., Schmitz, C.O., Parker, K.R., Woo, E., Chang, H.Y., Horvath, T.L., Carette, J.E., Bertozzi, C.R., Wilen, C.B., Satpathy, A.T., 2021. Discovery and functional interrogation of SARS-CoV-2 RNA-host protein interactions. Cell 184, 2394–2411.e16. https://doi.org/10.1016/j.cell.2021.03.012

Fraietta, J.A., Lacey, S.F., Orlando, E.J., Pruteanu-Malinici, I., Gohil, M., Lundh, S., Boesteanu, A.C., Wang, Y., O’Connor, R.S., Hwang, W.-T., Pequignot, E., Ambrose, D.E., Zhang, C., Wilcox, N., Bedoya, F., Dorfmeier, C., Chen, F., Tian, L., Parakandi, H., Gupta, M., Young, R.M., Johnson, F.B., Kulikovskaya, I., Liu, L., Xu, J., Kassim, S.H., Davis, M.M., Levine, B.L., Frey, N.V., Siegel, D.L., Huang, A.C., Wherry, E.J., Bitter, H., Brogdon, J.L., Porter, D.L., June, C.H., Melenhorst, J.J., 2018a. Determinants of response and resistance to CD19 chimeric antigen receptor (CAR) T cell therapy of chronic lymphocytic leukemia. Nat Med 24, 563–571. https://doi.org/10.1038/s41591-018-0010-1

Fraietta, J.A., Nobles, C.L., Sammons, M.A., Lundh, S., Carty, S.A., Reich, T.J., Cogdill, A.P., Morrissette, J.J.D., DeNizio, J.E., Reddy, S., Hwang, Y., Gohil, M., Kulikovskaya, I., Nazimuddin, F., Gupta, M., Chen, F., Everett, J.K., Alexander, K.A., Lin-Shiao, E., Gee, M.H., Liu, X., Young, R.M., Ambrose, D., Wang, Y., Xu, J., Jordan, M.S., Marcucci, K.T., Levine, B.L., Garcia, K.C., Zhao, Y., Kalos, M., Porter, D.L., Kohli, R.M., Lacey, S.F., Berger, S.L., Bushman, F.D., June, C.H., Melenhorst, J.J., 2018b. Disruption of TET2 promotes the therapeutic efficacy of CD19-targeted T cells. Nature 558, 307–312. https://doi.org/10.1038/s41586-018-0178-z

Gilbert, L.A., Horlbeck, M.A., Adamson, B., Villalta, J.E., Chen, Y., Whitehead, E.H., Guimaraes, C., Panning, B., Ploegh, H.L., Bassik, M.C., Qi, L.S., Kampmann, M., Weissman, J.S., 2014. Genome-Scale CRISPR-Mediated Control of Gene Repression and Activation. Cell 159, 647–661. https://doi.org/10.1016/j.cell.2014.09.029

Hao, Y., Hao, S., Andersen-Nissen, E., Mauck, W.M., Zheng, S., Butler, A., Lee, M.J., Wilk, A.J., Darby, C., Zager, M., Hoffman, P., Stoeckius, M., Papalexi, E., Mimitou, E.P., Jain, J., Srivastava, A., Stuart, T., Fleming, L.M., Yeung, B., Rogers, A.J., McElrath, J.M., Blish, C.A., Gottardo, R., Smibert, P., Satija, R., 2021. Integrated analysis of multimodal single-cell data. Cell 184, 3573–3587.e29. https://doi.org/10.1016/j.cell.2021.04.048

Hargreaves, D.C., Crabtree, G.R., 2011. ATP-dependent chromatin remodeling: genetics, genomics and mechanisms. Cell Res 21, 396–420. https://doi.org/10.1038/cr.2011.32

Henriksson, J., Chen, X., Gomes, T., Ullah, U., Meyer, K.B., Miragaia, R., Duddy, G., Pramanik, J., Yusa, K., Lahesmaa, R., Teichmann, S.A., 2019. Genome-wide CRISPR Screens in T Helper Cells Reveal Pervasive Crosstalk between Activation and Differentiation. Cell 176, 882–896.e18. https://doi.org/10.1016/j.cell.2018.11.044

Huang, H., Zhou, P., Wei, J., Long, L., Shi, H., Dhungana, Y., Chapman, N.M., Fu, G., Saravia, J., Raynor, J.L., Liu, S., Palacios, G., Wang, Y.-D., Qian, C., Yu, J., Chi, H., 2021. In vivo CRISPR screening reveals nutrient signaling processes underpinning CD8+ T cell fate decisions. Cell 184, 1245–1261.e21. https://doi.org/10.1016/j.cell.2021.02.021

Hudson, W.H., Gensheimer, J., Hashimoto, M., Wieland, A., Valanparambil, R.M., Li, P., Lin, J.-X., Konieczny, B.T., Im, S.J., Freeman, G.J., Leonard, W.J., Kissick, H.T., Ahmed, R., 2019. Proliferating Transitory T Cells with an Effector-like Transcriptional Signature Emerge from PD-1+ Stem-like CD8+ T Cells during Chronic Infection. Immunity 51, 1043–1058.e4. https://doi.org/10.1016/j.immuni.2019.11.002

Khan, O., Giles, J.R., McDonald, S., Manne, S., Ngiow, S.F., Patel, K.P., Werner, M.T., Huang, A.C., Alexander, K.A., Wu, J.E., Attanasio, J., Yan, P., George, S.M., Bengsch, B., Staupe, R.P., Donahue, G., Xu, W., Amaravadi, R.K., Xu, X., Karakousis, G.C., Mitchell, T.C., Schuchter, L.M., Kaye, J., Berger, S.L., Wherry, E.J., 2019. TOX transcriptionally and epigenetically programs CD8 + T cell exhaustion. Nature 571, 211–218. https://doi.org/10.1038/s41586-019-1325-x

Kim, D., Paggi, J.M., Park, C., Bennett, C., Salzberg, S.L., 2019. Graph-based genome alignment and genotyping with HISAT2 and HISAT-genotype. Nature Biotechnology 37, 907–915. https://doi.org/10.1038/s41587-019-0201-4

LaFleur, M.W., Nguyen, T.H., Coxe, M.A., Yates, K.B., Trombley, J.D., Weiss, S.A., Brown, F.D., Gillis, J.E., Coxe, D.J., Doench, J.G., Haining, W.N., Sharpe, A.H., 2019. A CRISPR-Cas9 delivery system for in vivo screening of genes in the immune system. Nature Communications 10, 1668. https://doi.org/10.1038/s41467-019-09656-2

Li, W., Xu, H., Xiao, T., Cong, L., Love, M.I., Zhang, F., Irizarry, R.A., Liu, J.S., Brown, M., Liu, X.S., 2014. MAGeCK enables robust identification of essential genes from genome-scale CRISPR/Cas9 knockout screens. Genome Biology 15, 554. https://doi.org/10.1186/s13059-014-0554-4

Long, A.H., Haso, W.M., Shern, J.F., Wanhainen, K.M., Murgai, M., Ingaramo, M., Smith, J.P., Walker, A.J., Kohler, M.E., Venkateshwara, V.R., Kaplan, R.N., Patterson, G.H., Fry, T.J., Orentas, R.J., Mackall, C.L., 2015. 4-1BB costimulation ameliorates T cell exhaustion induced by tonic signaling of chimeric antigen receptors. Nat. Med. 21, 581– 590. https://doi.org/10.1038/nm.3838

Love, M.I., Huber, W., Anders, S., 2014. Moderated estimation of fold change and dispersion for RNA-seq data with DESeq2. Genome Biology 15, 550. https://doi.org/10.1186/s13059-014-0550-8

Lynn, R.C., Weber, E.W., Sotillo, E., Gennert, D., Xu, P., Good, Z., Anbunathan, H., Lattin, J., Jones, R., Tieu, V., Nagaraja, S., Granja, J., de Bourcy, C.F.A., Majzner, R., Satpathy, A.T., Quake, S.R., Monje, M., Chang, H.Y., Mackall, C.L., 2019. c-Jun overexpression in CAR T cells induces exhaustion resistance. Nature 576, 293–300. https://doi.org/10.1038/s41586-019-1805-z

Mariathasan, S., Turley, S.J., Nickles, D., Castiglioni, A., Yuen, K., Wang, Y., Kadel Iii, E.E., Koeppen, H., Astarita, J.L., Cubas, R., Jhunjhunwala, S., Banchereau, R., Yang, Y., Guan, Y., Chalouni, C., Ziai, J., Şenbabaoğlu, Y., Santoro, S., Sheinson, D., Hung, J., Giltnane, J.M., Pierce, A.A., Mesh, K., Lianoglou, S., Riegler, J., Carano, R.A.D., Eriksson, P., Höglund, M., Somarriba, L., Halligan, D.L., van der Heijden, M.S., Loriot, Y., Rosenberg, J.E., Fong, L., Mellman, I., Chen, D.S., Green, M., Derleth, C., Fine, G.D., Hegde, P.S., Bourgon, R., Powles, T., 2018. TGFβ attenuates tumour response to PD-L1 blockade by contributing to exclusion of T cells. Nature 554, 544–548. https://doi.org/10.1038/nature25501

Mashtalir, N., D’Avino, A.R., Michel, B.C., Luo, J., Pan, J., Otto, J.E., Zullow, H.J., McKenzie, Z.M., Kubiak, R.L., St. Pierre, R., Valencia, A.M., Poynter, S.J., Cassel, S.H., Ranish, J.A., Kadoch, C., 2018. Modular Organization and Assembly of SWI/SNF Family Chromatin Remodeling Complexes. Cell 175, 1272–1288.e20. https://doi.org/10.1016/j.cell.2018.09.032

Mathur, R., Alver, B.H., San Roman, A.K., Wilson, B.G., Wang, X., Agoston, A.T., Park, P.J., Shivdasani, R.A., Roberts, C.W.M., 2017. ARID1A loss impairs enhancer-mediated gene regulation and drives colon cancer in mice. Nat Genet 49, 296–302. https://doi.org/10.1038/ng.3744

McKinney, E.F., Lee, J.C., Jayne, D.R.W., Lyons, P.A., Smith, K.G.C., 2015. T-cell exhaustion, co-stimulation and clinical outcome in autoimmunity and infection. Nature 523, 612–616. https://doi.org/10.1038/nature14468

McLane, L.M., Abdel-Hakeem, M.S., Wherry, E.J., 2019. CD8 T Cell Exhaustion During Chronic Viral Infection and Cancer. Annual Review of Immunology 37, 457–495. https://doi.org/10.1146/annurev-immunol-041015-055318

Miller, B.C., Sen, D.R., Al Abosy, R., Bi, K., Virkud, Y.V., LaFleur, M.W., Yates, K.B., Lako, A., Felt, K., Naik, G.S., Manos, M., Gjini, E., Kuchroo, J.R., Ishizuka, J.J., Collier, J.L., Griffin, G.K., Maleri, S., Comstock, D.E., Weiss, S.A., Brown, F.D., Panda, A., Zimmer, M.D., Manguso, R.T., Hodi, F.S., Rodig, S.J., Sharpe, A.H., Haining, W.N., 2019. Subsets of exhausted CD8 + T cells differentially mediate tumor control and respond to checkpoint blockade. Nature Immunology 20, 326–336. https://doi.org/10.1038/s41590-019-0312-6

Mondal, B., Jin, H., Kallappagoudar, S., Sedkov, Y., Martinez, T., Sentmanat, M.F., Poet, G.J., Li, C., Fan, Y., Pruett-Miller, S.M., Herz, H.-M., 2020. The histone deacetylase complex MiDAC regulates a neurodevelopmental gene expression program to control neurite outgrowth. Elife 9, e57519. https://doi.org/10.7554/eLife.57519

Morgens, D.W., Deans, R.M., Li, A., Bassik, M.C., 2016. Systematic comparison of CRISPR-Cas9 and RNAi screens for essential genes. Nat Biotechnol 34, 634–636. https://doi.org/10.1038/nbt.3567

Paley, M.A., Kroy, D.C., Odorizzi, P.M., Johnnidis, J.B., Dolfi, D.V., Barnett, B.E., Bikoff, E.K., Robertson, E.J., Lauer, G.M., Reiner, S.L., Wherry, E.J., 2012. Progenitor and Terminal Subsets of CD8+ T Cells Cooperate to Contain Chronic Viral Infection. Science 338, 1220–1225. https://doi.org/10.1126/science.1229620

Parnas, O., Jovanovic, M., Eisenhaure, T.M., Herbst, R.H., Dixit, A., Ye, C.J., Przybylski, D., Platt, R.J., Tirosh, I., Sanjana, N.E., Shalem, O., Satija, R., Raychowdhury, R., Mertins, P., Carr, S.A., Zhang, F., Hacohen, N., Regev, A., 2015. A Genome-wide CRISPR Screen in Primary Immune Cells to Dissect Regulatory Networks. Cell 162, 675–686. https://doi.org/10.1016/j.cell.2015.06.059

Pauken, K.E., Sammons, M.A., Odorizzi, P.M., Manne, S., Godec, J., Khan, O., Drake, A.M., Chen, Z., Sen, D.R., Kurachi, M., Barnitz, R.A., Bartman, C., Bengsch, B., Huang, A.C., Schenkel, J.M., Vahedi, G., Haining, W.N., Berger, S.L., Wherry, E.J., 2016. Epigenetic stability of exhausted T cells limits durability of reinvigoration by PD-1 blockade. Science 354, 1160–1165. https://doi.org/10.1126/science.aaf2807

Philip, M., Fairchild, L., Sun, L., Horste, E.L., Camara, S., Shakiba, M., Scott, A.C., Viale, A., Lauer, P., Merghoub, T., Hellmann, M.D., Wolchok, J.D., Leslie, C.S., Schietinger, A., 2017. Chromatin states define tumour-specific T cell dysfunction and reprogramming. Nature 545, 452–456. https://doi.org/10.1038/nature22367

Platt, R.J., Chen, S., Zhou, Y., Yim, M.J., Swiech, L., Kempton, H.R., Dahlman, J.E., Parnas, O., Eisenhaure, T.M., Jovanovic, M., Graham, D.B., Jhunjhunwala, S., Heidenreich, M., Xavier, R.J., Langer, R., Anderson, D.G., Hacohen, N., Regev, A., Feng, G., Sharp, P.A., Zhang, F., 2014. CRISPR-Cas9 Knockin Mice for Genome Editing and Cancer Modeling. Cell 159, 440–455. https://doi.org/10.1016/j.cell.2014.09.014

Pritykin, Y., van der Veeken, J., Pine, A.R., Zhong, Y., Sahin, M., Mazutis, L., Pe’er, D., Rudensky, A.Y., Leslie, C.S., 2021. A unified atlas of CD8 T cell dysfunctional states in cancer and infection. Mol Cell 81, 2477–2493.e10. https://doi.org/10.1016/j.molcel.2021.03.045

Raju, S., Xia, Y., Daniel, B., Yost, K.E., Bradshaw, E., Tonc, E., Verbaro, D.J., Kometani, K., Yokoyama, W.M., Kurosaki, T., Satpathy, A.T., Egawa, T., 2021. Identification of a T-bethi Quiescent Exhausted CD8 T Cell Subpopulation That Can Differentiate into TIM3+CX3CR1+ Effectors and Memory-like Cells. The Journal of Immunology. https://doi.org/10.4049/jimmunol.2001348

Replogle, J.M., Norman, T.M., Xu, A., Hussmann, J.A., Chen, J., Cogan, J.Z., Meer, E.J., Terry, J.M., Riordan, D.P., Srinivas, N., Fiddes, I.T., Arthur, J.G., Alvarado, L.J., Pfeiffer, K.A., Mikkelsen, T.S., Weissman, J.S., Adamson, B., 2020. Combinatorial single-cell CRISPR screens by direct guide RNA capture and targeted sequencing. Nat Biotechnol 38, 954– 961. https://doi.org/10.1038/s41587-020-0470-y

Ribas, A., Wolchok, J.D., 2018. Cancer immunotherapy using checkpoint blockade. Science 359, 1350–1355. https://doi.org/10.1126/science.aar4060

Roth, T.L., Li, P.J., Blaeschke, F., Nies, J.F., Apathy, R., Mowery, C., Yu, R., Nguyen, M.L.T., Lee, Y., Truong, A., Hiatt, J., Wu, D., Nguyen, D.N., Goodman, D., Bluestone, J.A., Ye, C.J., Roybal, K., Shifrut, E., Marson, A., 2020. Pooled Knockin Targeting for Genome Engineering of Cellular Immunotherapies. Cell 181, 728–744.e21. https://doi.org/10.1016/j.cell.2020.03.039

Sakuishi, K., Apetoh, L., Sullivan, J.M., Blazar, B.R., Kuchroo, V.K., Anderson, A.C., 2010. Targeting Tim-3 and PD-1 pathways to reverse T cell exhaustion and restore anti-tumor immunity. J Exp Med 207, 2187–2194. https://doi.org/10.1084/jem.20100643

Satpathy, A.T., Granja, J.M., Yost, K.E., Qi, Y., Meschi, F., McDermott, G.P., Olsen, B.N., Mumbach, M.R., Pierce, S.E., Corces, M.R., Shah, P., Bell, J.C., Jhutty, D., Nemec, C.M., Wang, J., Wang, L., Yin, Y., Giresi, P.G., Chang, A.L.S., Zheng, G.X.Y., Greenleaf, W.J., Chang, H.Y., 2019. Massively parallel single-cell chromatin landscapes of human immune cell development and intratumoral T cell exhaustion. Nat. Biotechnol. 37, 925–936. https://doi.org/10.1038/s41587-019-0206-z

Schep, A.N., Wu, B., Buenrostro, J.D., Greenleaf, W.J., 2017. chromVAR: inferring transcription-factor-associated accessibility from single-cell epigenomic data. Nat Methods 14, 975–978. https://doi.org/10.1038/nmeth.4401

Schietinger, A., Philip, M., Krisnawan, V.E., Chiu, E.Y., Delrow, J.J., Basom, R.S., Lauer, P., Brockstedt, D.G., Knoblaugh, S.E., Hämmerling, G.J., Schell, T.D., Garbi, N., Greenberg, P.D., 2016. Tumor-Specific T Cell Dysfunction Is a Dynamic Antigen-Driven Differentiation Program Initiated Early during Tumorigenesis. Immunity 45, 389–401. https://doi.org/10.1016/j.immuni.2016.07.011

Scott, A.C., Dündar, F., Zumbo, P., Chandran, S.S., Klebanoff, C.A., Shakiba, M., Trivedi, P., Menocal, L., Appleby, H., Camara, S., Zamarin, D., Walther, T., Snyder, A., Femia, M.R., Comen, E.A., Wen, H.Y., Hellmann, M.D., Anandasabapathy, N., Liu, Y., Altorki, N.K., Lauer, P., Levy, O., Glickman, M.S., Kaye, J., Betel, D., Philip, M., Schietinger, A., 2019. TOX is a critical regulator of tumour-specific T cell differentiation. Nature 571, 270–274. https://doi.org/10.1038/s41586-019-1324-y

Scott-Browne, J.P., López-Moyado, I.F., Trifari, S., Wong, V., Chavez, L., Rao, A., Pereira, R.M., 2016. Dynamic Changes in Chromatin Accessibility Occur in CD8+ T Cells Responding to Viral Infection. Immunity 45, 1327–1340. https://doi.org/10.1016/j.immuni.2016.10.028

Sen, D.R., Kaminski, J., Barnitz, R.A., Kurachi, M., Gerdemann, U., Yates, K.B., Tsao, H.-W., Godec, J., LaFleur, M.W., Brown, F.D., Tonnerre, P., Chung, R.T., Tully, D.C., Allen, T.M., Frahm, N., Lauer, G.M., Wherry, E.J., Yosef, N., Haining, W.N., 2016. The epigenetic landscape of T cell exhaustion. Science 354, 1165–1169. https://doi.org/10.1126/science.aae0491

Seo, H., González-Avalos, E., Zhang, W., Ramchandani, P., Yang, C., Lio, C.-W.J., Rao, A., Hogan, P.G., 2021. BATF and IRF4 cooperate to counter exhaustion in tumor-infiltrating CAR T cells. Nat Immunol 1–13. https://doi.org/10.1038/s41590-021-00964-8

Shalem, O., Sanjana, N.E., Hartenian, E., Shi, X., Scott, D.A., Mikkelson, T., Heckl, D., Ebert, B.L., Root, D.E., Doench, J.G., Zhang, F., 2014. Genome-scale CRISPR-Cas9 knockout screening in human cells. Science 343, 84–87. https://doi.org/10.1126/science.1247005

Shannon, P., Markiel, A., Ozier, O., Baliga, N.S., Wang, J.T., Ramage, D., Amin, N., Schwikowski, B., Ideker, T., 2003. Cytoscape: a software environment for integrated models of biomolecular interaction networks. Genome Res 13, 2498–2504. https://doi.org/10.1101/gr.1239303

Shifrut, E., Carnevale, J., Tobin, V., Roth, T.L., Woo, J.M., Bui, C.T., Li, P.J., Diolaiti, M.E., Ashworth, A., Marson, A., 2018. Genome-wide CRISPR Screens in Primary Human T Cells Reveal Key Regulators of Immune Function. Cell 175, 1958–1971.e15. https://doi.org/10.1016/j.cell.2018.10.024

Singer, M., Wang, C., Cong, L., Marjanovic, N.D., Kowalczyk, M.S., Zhang, H., Nyman, J., Sakuishi, K., Kurtulus, S., Gennert, D., Xia, J., Kwon, J.Y.H., Nevin, J., Herbst, R.H., Yanai, I., Rozenblatt-Rosen, O., Kuchroo, V.K., Regev, A., Anderson, A.C., 2016. A Distinct Gene Module for Dysfunction Uncoupled from Activation in Tumor-Infiltrating T Cells. Cell 166, 1500–1511.e9. https://doi.org/10.1016/j.cell.2016.08.052

Spitzer, M.H., Carmi, Y., Reticker-Flynn, N.E., Kwek, S.S., Madhireddy, D., Martins, M.M., Gherardini, P.F., Prestwood, T.R., Chabon, J., Bendall, S.C., Fong, L., Nolan, G.P., Engleman, E.G., 2017. Systemic Immunity Is Required for Effective Cancer Immunotherapy. Cell 168, 487–502.e15. https://doi.org/10.1016/j.cell.2016.12.022

Stadtmauer, E.A., Fraietta, J.A., Davis, M.M., Cohen, A.D., Weber, K.L., Lancaster, E., Mangan, P.A., Kulikovskaya, I., Gupta, M., Chen, F., Tian, L., Gonzalez, V.E., Xu, J., Jung, I., Melenhorst, J.J., Plesa, G., Shea, J., Matlawski, T., Cervini, A., Gaymon, A.L., Desjardins, S., Lamontagne, A., Salas-Mckee, J., Fesnak, A., Siegel, D.L., Levine, B.L., Jadlowsky, J.K., Young, R.M., Chew, A., Hwang, W.-T., Hexner, E.O., Carreno, B.M., Nobles, C.L., Bushman, F.D., Parker, K.R., Qi, Y., Satpathy, A.T., Chang, H.Y., Zhao, Y., Lacey, S.F., June, C.H., 2020. CRISPR-engineered T cells in patients with refractory cancer. Science 367. https://doi.org/10.1126/science.aba7365

Szklarczyk, D., Gable, A.L., Lyon, D., Junge, A., Wyder, S., Huerta-Cepas, J., Simonovic, M., Doncheva, N.T., Morris, J.H., Bork, P., Jensen, L.J., Mering, C. von, 2019. STRING v11: protein-protein association networks with increased coverage, supporting functional discovery in genome-wide experimental datasets. Nucleic Acids Res 47, D607–D613. https://doi.org/10.1093/nar/gky1131

Vardhana, S.A., Hwee, M.A., Berisa, M., Wells, D.K., Yost, K.E., King, B., Smith, M., Herrera, P.S., Chang, H.Y., Satpathy, A.T., van den Brink, M.R.M., Cross, J.R., Thompson, C.B., 2020. Impaired mitochondrial oxidative phosphorylation limits the self-renewal of T cells exposed to persistent antigen. Nature Immunology 21, 1022–1033. https://doi.org/10.1038/s41590-020-0725-2

Vierbuchen, T., Ling, E., Cowley, C.J., Couch, C.H., Wang, X., Harmin, D.A., Roberts, C.W.M., Greenberg, M.E., 2017. AP-1 Transcription Factors and the BAF Complex Mediate Signal-Dependent Enhancer Selection. Mol Cell 68, 1067–1082.e12. https://doi.org/10.1016/j.molcel.2017.11.026

Wang, T., Wei, J.J., Sabatini, D.M., Lander, E.S., 2014. Genetic Screens in Human Cells Using the CRISPR-Cas9 System. Science 343, 80–84. https://doi.org/10.1126/science.1246981

Weber, E.W., Parker, K.R., Sotillo, E., Lynn, R.C., Anbunathan, H., Lattin, J., Good, Z., Belk, J.A., Daniel, B., Klysz, D., Malipatlolla, M., Xu, P., Bashti, M., Heitzeneder, S., Labanieh, L., Vandris, P., Majzner, R.G., Qi, Y., Sandor, K., Chen, L.-C., Prabhu, S., Gentles, A.J., Wandless, T.J., Satpathy, A.T., Chang, H.Y., Mackall, C.L., 2021. Transient rest restores functionality in exhausted CAR-T cells through epigenetic remodeling. Science 372. https://doi.org/10.1126/science.aba1786

Wei, J., Long, L., Zheng, W., Dhungana, Y., Lim, S.A., Guy, C., Wang, Y., Wang, Y.-D., Qian, C., Xu, B., Kc, A., Saravia, J., Huang, H., Yu, J., Doench, J.G., Geiger, T.L., Chi, H., 2019. Targeting REGNASE-1 programs long-lived effector T cells for cancer therapy. Nature 576, 471–476. https://doi.org/10.1038/s41586-019-1821-z

Wei, S.C., Levine, J.H., Cogdill, A.P., Zhao, Y., Anang, N.-A.A.S., Andrews, M.C., Sharma, P., Wang, J., Wargo, J.A., Pe’er, D., Allison, J.P., 2017. Distinct Cellular Mechanisms Underlie Anti-CTLA-4 and Anti-PD-1 Checkpoint Blockade. Cell 170, 1120–1133.e17. https://doi.org/10.1016/j.cell.2017.07.024

Wherry, E.J., Kurachi, M., 2015. Molecular and cellular insights into T cell exhaustion. Nat Rev Immunol 15, 486–499. https://doi.org/10.1038/nri3862

Xu, G., Chhangawala, S., Cocco, E., Razavi, P., Cai, Y., Otto, J.E., Ferrando, L., Selenica, P., Ladewig, E., Chan, C., Da Cruz Paula, A., Witkin, M., Cheng, Y., Park, J., Serna-Tamayo, C., Zhao, H., Wu, F., Sallaku, M., Qu, X., Zhao, A., Collings, C.K., D’Avino, A.R., Jhaveri, K., Koche, R., Levine, R.L., Reis-Filho, J.S., Kadoch, C., Scaltriti, M., Leslie, C.S., Baselga, J., Toska, E., 2020. ARID1A determines luminal identity and therapeutic response in estrogen-receptor-positive breast cancer. Nat Genet 52, 198– 207. https://doi.org/10.1038/s41588-019-0554-0

Yao, W., King, D.A., Beckwith, S.L., Gowans, G.J., Yen, K., Zhou, C., Morrison, A.J., 2016. The INO80 Complex Requires the Arp5-Ies6 Subcomplex for Chromatin Remodeling and Metabolic Regulation. Mol Cell Biol 36, 979–991. https://doi.org/10.1128/MCB.00801-15

Yost, K.E., Chang, H.Y., Satpathy, A.T., 2021. Recruiting T cells in cancer immunotherapy. Science 372, 130–131. https://doi.org/10.1126/science.abd1329

Yost, K.E., Satpathy, A.T., Wells, D.K., Qi, Y., Wang, C., Kageyama, R., McNamara, K.L., Granja, J.M., Sarin, K.Y., Brown, R.A., Gupta, R.K., Curtis, C., Bucktrout, S.L., Davis, M.M., Chang, A.L.S., Chang, H.Y., 2019. Clonal replacement of tumor-specific T cells following PD-1 blockade. Nat. Med. 25, 1251–1259. https://doi.org/10.1038/s41591-019-0522-3

Zajac, A.J., Blattman, J.N., Murali-Krishna, K., Sourdive, D.J., Suresh, M., Altman, J.D., Ahmed, R., 1998. Viral immune evasion due to persistence of activated T cells without effector function. J. Exp. Med. 188, 2205–2213. https://doi.org/10.1084/jem.188.12.2205

Zander, R., Schauder, D., Xin, G., Nguyen, C., Wu, X., Zajac, A., Cui, W., 2019. CD4+ T Cell Help Is Required for the Formation of a Cytolytic CD8+ T Cell Subset that Protects against Chronic Infection and Cancer. Immunity 51, 1028–1042.e4. https://doi.org/10.1016/j.immuni.2019.10.009

Zhang, Y., Liu, T., Meyer, C.A., Eeckhoute, J., Johnson, D.S., Bernstein, B.E., Nusbaum, C., Myers, R.M., Brown, M., Li, W., Liu, X.S., 2008. Model-based Analysis of ChIP-Seq (MACS). Genome Biol 9, R137. https://doi.org/10.1186/gb-2008-9-9-r137

